# Modulation of social behavior in adult zebrafish (*Danio rerio*): a systematic review and meta-analysis

**DOI:** 10.64898/2026.05.28.728488

**Authors:** Daniela V. Müller, Matheus Gallas-Lopes, Mariana B. Abreu, Bruno D. Arbo, Leonardo M. Bastos, Nicole T. Fröhlich, Matheus Marcon, Izabela B. Moraes, Luiz Cesar C. P. da Silva, Gabriel da R. Zurchimitten, Ana P. Herrmann

## Abstract

Social behavior is a fundamental phenotype across vertebrates. Zebrafish (*Danio rerio*) have emerged as a valuable translational model for investigating the neurobiological mechanisms underlying sociability, particularly due to their robust shoaling behavior and experimental tractability. However, the literature presents issues of reproducibility and inconsistent findings regarding the modulation of social preference and shoal cohesion in adult zebrafish. We conducted a systematic review and meta-analysis to synthesize studies evaluating the effects of pharmacological interventions that modulate the central nervous system and stress-related interventions on social behavior in adult zebrafish and, when available, anxiety-like behavior. The literature search was performed in three databases (Embase, PubMed, and Web of Science), followed by a two-step screening process based on inclusion/exclusion criteria. The included studies underwent extraction of qualitative and quantitative data, as well as risk of bias assessment. Interventions from the included studies (n = 108) were categorized according to their nature, mechanism of action, and/or therapeutic purpose, resulting in seven, four, and five meta-analyses for social preference, shoal cohesion, and anxiety-related tests, respectively. Ethanol, NMDA antagonists, pro-dopaminergic agents, and stress-related interventions decreased social preference, while stress-related interventions increased shoal cohesion. The fact that stress produced opposite effects suggests that these paradigms measure distinct sociability constructs, or perhaps are differentially modulated by confounding factors, like anxiety for example. The studies presented high heterogeneity, with prediction intervals compatible with effects in both directions, as well as methodological limitations and deficiencies in data reporting, as evidenced by the risk of bias assessment. These findings emphasize the need for well-designed new studies to validate the findings and expand the evidence on interventions that currently lack sufficient studies for quantitative synthesis.

## 1. Introduction

Social behavior is a fundamental phenotype for the survival and reproduction of vertebrate species (Chen & Hong, 2018; Robinson et al., 2019; Wei et al., 2021). Characterizing social interaction in both humans and animals is essential for advancing our understanding of the mechanisms and neurobiological pathways underlying social behavior (Hofmann et al., 2014; Saverino & Gerlai, 2008). In this context, animal models have been extensively used to investigate these processes and their disruption, providing a promising approach for studying a phenotype with relevance to neuropsychiatric disorders such as autism, schizophrenia, and depression (Henry et al., 2016; Lieberman et al., 2001; Marotta et al., 2020; Soares et al., 2018).

Notably, the neural circuits governing social behavior are highly conserved across species, enabling the use of alternative model organisms in translational research (Geng & Peterson, 2019; Goodson, 2005; Newman, 1999). Among these organisms, the zebrafish (*Danio rerio,* Hamilton, 1822) has become prominent in neuroscience research, particularly for investigating social deficits related to human conditions (Kalueff et al., 2014; Lieschke & Currie, 2007; Norton, 2013; Stewart et al., 2014; Tropepe & Sive, 2003). A key feature of zebrafish sociality is shoaling behavior, defined as the innate tendency to swim in close proximity to conspecifics (Spence et al., 2008; Suriyampola et al., 2016). This behavior plays a crucial role in the species’ ecology, conferring adaptive benefits such as enhanced predator avoidance, optimized foraging, and hydrodynamic efficiency. Importantly, shoal cohesion is modulated by a range of environmental and intrinsic factors, including individual variability and physiological states such as stress and nutritional deprivation (Maaswinkel et al., 2013b; Miller & Gerlai, 2007; Qin et al., 2014; Ruhl et al., 2009; Spence et al., 2008; Wright et al., 2003, 2006).

Under laboratory conditions, zebrafish social behavior is quantitatively assessed primarily using two well-established assays: the social preference and the shoal cohesion tests (Apukhtin et al., 2026; Fontana et al., 2022). A growing body of research has examined the effects of diverse interventions on zebrafish social behavior, including pharmacological agents (e.g., anxiolytics and antidepressants), stress-related manipulations, and experimental models relevant to neuropsychiatric disorders (Gebauer et al., 2011; Giacomini et al., 2016; Lee et al., 2010; Miller et al., 2013; Savio et al., 2012; Schaefer et al., 2015). However, this literature is marked by substantial methodological heterogeneity, inconsistent findings, and discrepancies in the interpretation of results, challenging the reproducibility and translational relevance of the assays (Avey et al., 2016; Gosselin, 2021; Sison et al., 2006).

A systematic review with meta-analysis is thus warranted to integrate the available evidence, identify sources of variability, and clarify the interpretation of findings on zebrafish social behavior. This will also inform future experimental designs and promote greater methodological rigor in the field. We conducted a systematic review to address the following PICO question: what are the effects of different pharmacological and stress-related interventions (intervention) on social behavior (outcome) in adult zebrafish (population)? We also evaluated anxiety-related behavior in the included studies to aid the interpretation of sociability measures.

## 2. Methods

The reporting of this study followed the Preferred Reporting Items for Systematic Reviews and Meta-Analyses (PRISMA) guidelines (Page et al., 2021). All data and materials associated with this study, including extracted datasets and analysis code, are publicly available via OSF at <u>osf.io/h4v3k</u> (Gallas-Lopes et al., 2024) and GitHub at <u>github.com/BRISAcollab/social-zebrafish/</u> (BRISA, 2026).

### 2.1. Protocol registration

The protocol for this study was developed in accordance with the SYRCLE guidelines for systematic reviews of animal intervention studies (version 2.0, December 2014) (de Vries et al., 2015) and deposited in the Open Science Framework (OSF) on January 8, 2025. The preregistration is publicly available at https://osf.io/x95w6 (Gallas-Lopes et al., 2025).

The aim of the study was to evaluate and summarize the effects of a selected set of pharmacological and stress-related interventions on zebrafish social behavior, also investigating whether anxiety-like behavior is modulated in parallel by the same interventions.

### 2.2. Search strategy and eligibility criteria

A literature search was performed in three bibliographic databases: Embase, PubMed, and Web of Science. Search strategies combined terms for the population (zebrafish) and the outcome of interest (social behavior), with adaptations to each database. The complete query for each database is available at <u>osf.io/x95w6/files/f8p7w</u>. The search encompassed all articles indexed up to October 25^th^, 2024, with no restrictions on publication language. The bibliographic data from the databases were imported into Rayyan platform (Rayyan Systems Inc, Cambridge, MA, USA, https://www.rayyan.ai) (Ouzzani et al., 2016), where duplicates were detected and removed. After duplicate removal, eligible studies were screened by title and abstract, followed by full-text assessment. Two reviewers independently screened each record (D.V.M., M.G.L., M.B.A., B.D.A., L.B.M., N.T.F., I.B.M., L.C.C.P.S., G.R.Z., A.P.H.). At both screening stages, studies were excluded only upon consensus between two reviewers, and any disagreements at the full-text stage were resolved through discussion between the reviewers or consultation with the other team members. At the title and abstract stage, studies were assessed for eligibility based solely on type of record or population. If at least one reviewer marked a record as “maybe”, the study moved to full-text screening. At the full-text stage, studies were included if they met all of the following criteria:

1. Type of record: peer-reviewed articles or preprints.
2. Population: adult zebrafish at the time the outcome was assessed.
3. Intervention: pharmacological treatments that modulate the central nervous system or stress-related interventions that were applied directly to the animals.
4. Comparator: control group not exposed to the intervention.
5. Outcome measures: social preference or shoal cohesion tests.
6. Study design: controlled studies with separate treatment arms.
7. Data availability: quantitative data (mean, standard deviation (SD) or standard error of the mean (SEM), and sample size) reported for the outcomes of interest.

The inclusion and exclusion guide applied during the screening phase are available in the OSF preregistration (https://osf.io/h4v3k/files/6ahzg).

### 2.3. Data extraction

Qualitative and quantitative data were collected independently by two investigators using custom-designed forms on the Jotform online platform (Jotform Inc, San Francisco, CA, USA, https://www.jotform.com). All authors contributed to data extraction. Any discrepancies in the extracted data were resolved by a third investigator (D.V.M.). The following qualitative data were extracted from the included studies: study identification (first author’s last name and publication year); strain, sex, age, and developmental stage at the time of intervention; intervention type (pharmacological or stress-related), name of the intervention, administration route, dose, schedule (acute or repeated/continuous), and duration. The qualitative extraction guide is available at OSF (https://osf.io/h4v3k/files/2bsn5), and the raw qualitative data independently extracted by the two investigators are also publicly available at OSF (https://osf.io/h4v3k/files/3qgd6).

A single study could contribute data from multiple experiments and multiple comparisons of interest. Experiments were defined as independent samples of animals, either because they differed in population or protocol (e.g., sex, age, strain, developmental stage, timing of outcome assessment), or because they represented separate experimental runs using different sets of animals. Each experiment was treated as a separate experimental unit for data extraction. Within a given experiment, multiple intervention groups could be compared with a shared control group. All intervention-control comparisons of interest were extracted. Thus, data were structured hierarchically, with comparisons nested within experiments, and experiments nested within studies.

All behavioral outcomes related to social behavior in adult zebrafish following interventions affecting the central nervous system (and measured by the social preference or shoal cohesion tests) were extracted. In this review, social preference was defined as the interest or proximity displayed by an individual zebrafish toward a conspecific with restricted movement, whereas shoal cohesion was defined as direct social interaction among freely moving conspecifics forming a shoal. Studies using different terminologies but assessing comparable behavioral constructs were categorized according to these operational definitions. In addition, anxiety-like measures obtained from the novel tank, light-dark, and/or open tank tests were extracted whenever available, regardless of whether they were assessed in the same animal cohort as the social behavior tests. When a single study reported anxiety-related outcomes from multiple assays, it was frequently unclear whether different cohorts were used or the same animals were subjected to sequential testing. Therefore, we could not precisely account for within-subject correlations. To prevent the uneven application of an additional correction layer for the few studies where sequential testing was explicit, non-independence corrections were thus limited to shared control groups and within-study experiments.

When different measures were reported for a given behavioral assay, a frequency-based hierarchy was applied to select a single measure per test. When multiple measures based on time (e.g., absolute time and percent time) were reported, the absolute values were preferred. When the same animal cohort was assessed more than once in the same test, only the first assessment was extracted.

When multiple data points from a given test were reported only as time-course data and no summary measure was provided, only the first time point was extracted.

Whenever available, the values for the quantitative synthesis were extracted from text or tables. Otherwise, WebPlotDigitizer (v4.5, Automeris LLC, Pacifica, CA, USA, https://automeris.io) was used to manually estimate the values from the graphs. The mean, standard deviation (SD) or standard error of the mean (SEM), and sample size (n) were extracted for all groups of interest. If the study reported the SEM, SD was calculated by multiplying the SEM by the square root of the sample size. When the sample size was reported as a range, the lower bound was used to compute effect sizes, whereas the upper bound was used to calculate the corresponding SD. The quantitative extraction guide is available at OSF (https://osf.io/h4v3k/files/cvfjy), and the raw quantitative data independently extracted by the two investigators are also publicly available at OSF (https://osf.io/h4v3k/files/vhw7f).

### 2.4. Risk of bias assessment

The risk of bias of each study was assessed by at least two independent reviewers (all authors contributed to this assessment). Only the relevant experiments and outcomes were considered for the judgment, and the scores were decided per study. Any discrepancies were resolved through discussion among the reviewers or consultation with the other team members to reach a consensus. The criteria were based on the SYRCLE’s Risk of Bias tool (Hooijmans et al., 2014) with the following adaptations: item 3 (allocation concealment) was omitted, and item 10 (other sources of bias) was specified to address pseudoreplication (treating non-independent observations as independent experimental units in the analysis) and lack of procedural equivalence (not exposing control groups to the same procedures as intervention groups, except for the active treatment).

We thus evaluated the following items to assess methodological quality: sequence generation (description of random allocation of animals to experimental groups); baseline characteristics; random housing conditions during the experiments; blinding of caregivers and/or investigators during the experiment; random selection for outcome assessment; blinding of investigators during outcome assessment; incomplete outcome data; selective outcome reporting; pseudoreplication; procedural equivalence. Risk of bias was rated as “low” when appropriate measures to minimize bias were implemented and clearly reported. Studies were rated as “high” when bias-control measures were not implemented (e.g., lack of blinding, non-random selection for outcome assessment) or when reporting inconsistencies were detected (e.g., unexplained missing data, specified outcome not reported). When reporting was insufficient to judge, items were rated as “unclear.” Studies with two or more “high” ratings were classified as having a high overall risk of bias, while the remaining studies were classified as low overall risk; this classification was used in sensitivity analyses.

We additionally annotated whether blinding of the investigators to group allocation was impossible due to the nature of the intervention (e.g., application of a physical stressor), and whether the study reported a preregistered protocol. Both items were classified as “yes” or “no.” The risk of bias assessment guide is available at OSF (https://osf.io/h4v3k/files/6dma8), and the raw data independently extracted by the two investigators are also publicly available at OSF (https://osf.io/h4v3k/files/st2kr).

### 2.5. Meta-analyses

Interventions were categorized according to their nature, mechanism of action, and/or therapeutic purpose. Separate meta-analyses were conducted for the primary outcomes of social preference and shoal cohesion when at least five studies were available for a given intervention category. Meta-analyses for the secondary outcome of anxiety-like behavior were conducted using the same criterion. For interventions that did not meet the minimum study threshold, effect sizes were nevertheless computed and grouped under a “miscellaneous” category, but these data were not meta-analyzed (see Supplementary Material).

When pseudoreplication was detected, sample sizes were adjusted to reflect the true number of independent experimental units (e.g., shoals rather than sequential screenshots), preventing the artificial inflation of study precision in the meta-analysis. Effect sizes were reversed (multiplied by -1) when necessary to standardize the direction of effects for specific behavioral traits, ensuring consistent interpretation of intervention effects across social behavior and anxiety-related outcomes.

To account for the non-independence of comparisons extracted from the same study and/or experiment, a multilevel and multivariate meta-analysis was conducted. Effect sizes were calculated as standardized mean differences (SMDs) using Hedges’ g method. Correlated hierarchical random-effects models were fitted with random intercepts for study and experiment, and a within-experiment correlation coefficient (⍴) of 0.5 was assumed. Models were fitted using restricted maximum likelihood estimation, and cluster-robust variance estimation at the study level was applied using the clubSandwich package (Pustejovsky et al., 2025) to provide additional protection against residual dependence. Heterogeneity was assessed using Cochran’s Q (significance set at p < 0.1), total I², and σ² at each random-effects level. Prediction intervals were estimated to reflect the expected range of true effects in future studies, accounting for between-study heterogeneity. To investigate potential sources of heterogeneity, we conducted a meta-regression with ethanol concentration included as a moderating variable. Analyses of additional variables as potential moderators were not conducted due to insufficient number of studies to support robust testing. Meta-analyses were performed in R (R Core Team 2023) using the packages “metafor” (Viechtbauer, 2010) and “orchaRd” (Nakagawa et al., 2023).

### 2.6. Small-study effects

We assessed small-study effects for each meta-analysis through (a) visual inspection of the funnel plots and (b) Egger-type regression tests. Funnel plots were visually inspected to evaluate asymmetry in the distribution of effect sizes in relation to study precision. We examined the association between effect sizes and precision using two regression-based approaches. In the first approach, precision was represented by the standard error obtained from the sampling variance of the SMDs. In the second approach, precision was represented by an alternative “effective” standard error derived solely from the intervention and control group sample sizes, thereby reducing the direct dependence between the effect size estimate and its standard error (Nakagawa et al., 2022). Evidence of small-study effects and potential publication bias was considered when asymmetry was observed in the funnel plots and when the intercept from the effect size by precision regression was significantly different from zero (p < 0.05).

### 2.7. Sensitivity analysis

Sensitivity analyses were conducted to assess the robustness of the main findings using three approaches. These included (a) a leave-one-study-out procedure, in which the meta-analysis was iteratively repeated after excluding one study at a time to identify potentially influential studies; (b) exclusion of studies rated as having a high overall risk of bias to evaluate whether results were driven by lower-quality evidence; and (c) variation of the within-study correlation coefficient (⍴ = 0.0, 0.2, 0.4, 0.5, 0.6, 0.8) used to impute the covariance matrix, to assess whether pooled estimates remained stable under different assumptions about outcome dependency.

### 2.8. Protocol deviations

According to the original protocol, all measures of social preference and/or shoal cohesion, as well as anxiety-like behavior outcomes, were planned to be extracted. During the extraction process, however, it became evident that studies frequently reported multiple outcome measures for the same animals. To limit further data non-independence, we deviated from the protocol by implementing a hierarchical selection strategy based on outcome frequency.

The protocol originally planned a secondary analysis considering animal groups subjected to a disease model and exposed to the intervention of interest. However, this analysis was not conducted because of an insufficient number of studies.

According to the protocol, shared control groups would be accounted for by dividing their sample size by the number of multiple comparisons. However, we refined the analysis and instead applied a statistical model that explicitly accounts for codependencies among effect sizes through the use of a variance-covariance matrix. This approach allowed us to retain the original sample sizes while appropriately modeling the shared structure among comparisons.

## 3. Results

### 3.1. Study characteristics

Our search retrieved a total of 3,177 records. After the removal of duplicates, 1,719 records were screened based on titles and abstracts. Following the exclusion of records that did not meet the inclusion criteria for record type or population, 1,131 studies were assessed for eligibility based on full-text screening. Of these, 108 studies met the inclusion criteria and were included in the final analysis (Figure 1). The complete list of included studies is available at https://osf.io/h4v3k/files/hzwe9.

**Figure 1.**
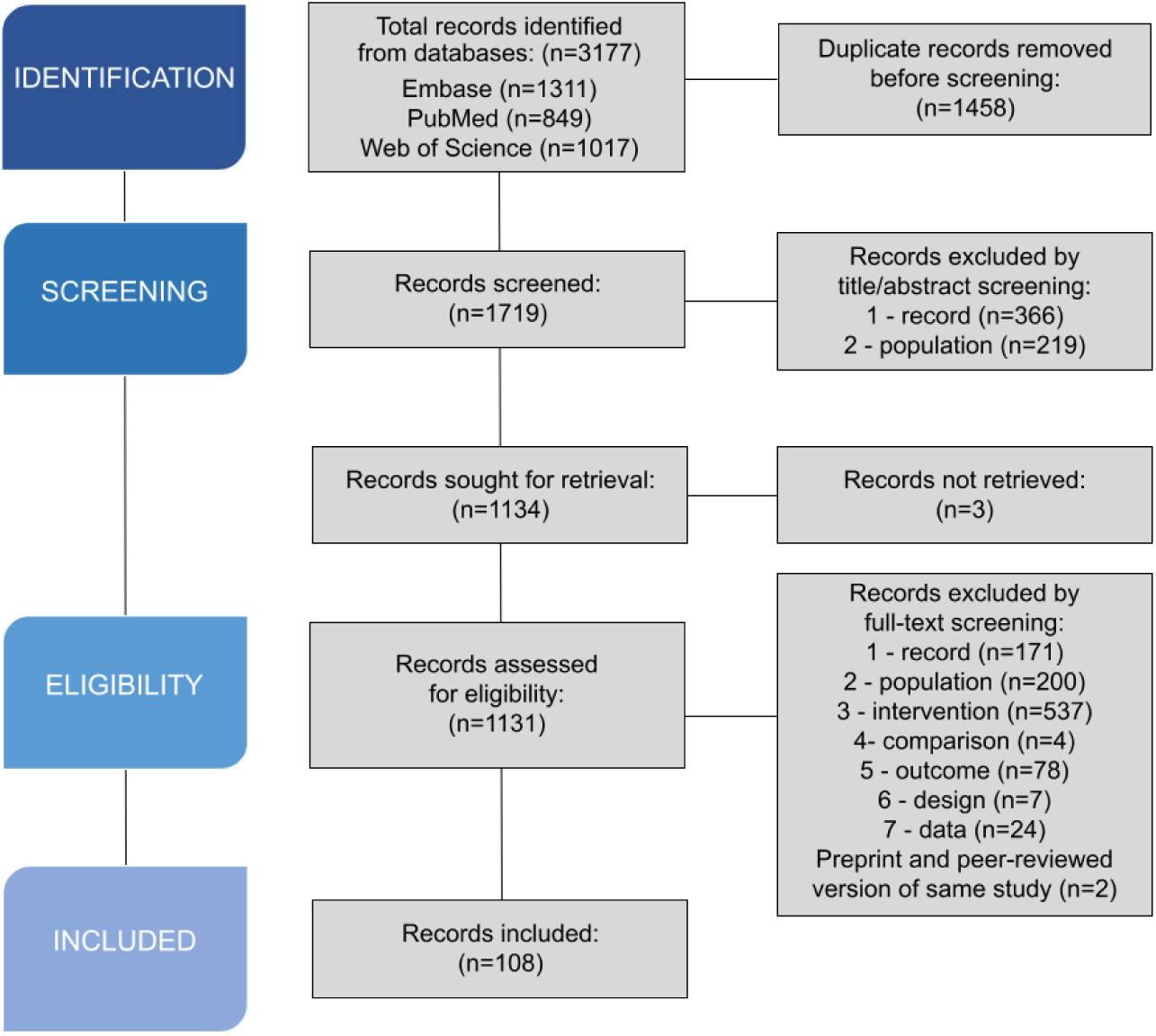
Flowchart diagram of the collection of studies and selection process.

Among the 108 included studies, 65 provided data from the social preference test, 30 from the shoal cohesion test, and 13 from both tests. Most studies used mixed-sex groups (62.96%, 68), whereas 13.89% (15) used only males and 7.41% (8) only females; sex was unclear in 16.67% (18) of studies. The qualitative and quantitative raw data on social behavior for each study are available in Supplementary Table S1. Furthermore, 63 studies evaluated anxiety-like behavioral outcomes using the novel tank (57), light-dark (18), and/or open tank (12) tests. The corresponding data on anxiety-like behavior are provided in Supplementary Table S2.

### 3.2. Risk of bias

As shown in Figure 2, insufficient reporting resulted in unclear judgments for random sequence generation in 87% of studies, baseline comparability between experimental groups in 65%, and random housing allocation in 97%. Regarding investigator blinding, 19% of studies were classified as high risk of bias; however, in all but two (1.85%) of these cases blinding was inherently unfeasible due to the nature of the intervention (e.g., chronic stress protocols). In the studies in which blinding was possible, it was reported in only 9.26% (10) of them. Bias related to random outcome assessment and blinding during outcome assessment was rated as unclear in 79% and 72% of studies, respectively. It should be noted that studies only reporting the use of automated software for outcome assessment were rated as unclear. Although blinding during outcome assessment is readily feasible, two studies reported that this measure was not implemented. For incomplete outcome data, 50% of studies were rated as unclear risk of bias due to poor reporting that prevented cross-checking allocated animals and those included in the analysis, while 20% were rated as high risk because of unexplained inconsistencies in reported sample sizes or degrees of freedom in the statistical analyses. Selective outcome reporting was rated as low risk of bias in 81% of studies, as preregistration was not required in our assessment, and only outcomes related to social behavior were cross-checked within the article. Nevertheless, 9% of studies mentioned outcomes that were not subsequently reported. Pseudoreplication was identified as a major source of bias, with 44% of studies classified as high risk for this item. This issue was particularly frequent in shoal cohesion assays, where some studies inappropriately treated sequential video frames or screen captures of the same shoal as independent experimental units. Whenever possible, sample sizes in the meta-analyses were corrected for inflation arising from pseudoreplication; however, this was not always feasible due to poor reporting. Considering procedural equivalence, 19% of studies were rated as high risk of bias, largely because procedural equivalence was not achievable for certain interventions given their nature. None of the studies included in this systematic review mentioned a preregistered study protocol.

**Figure 2.**
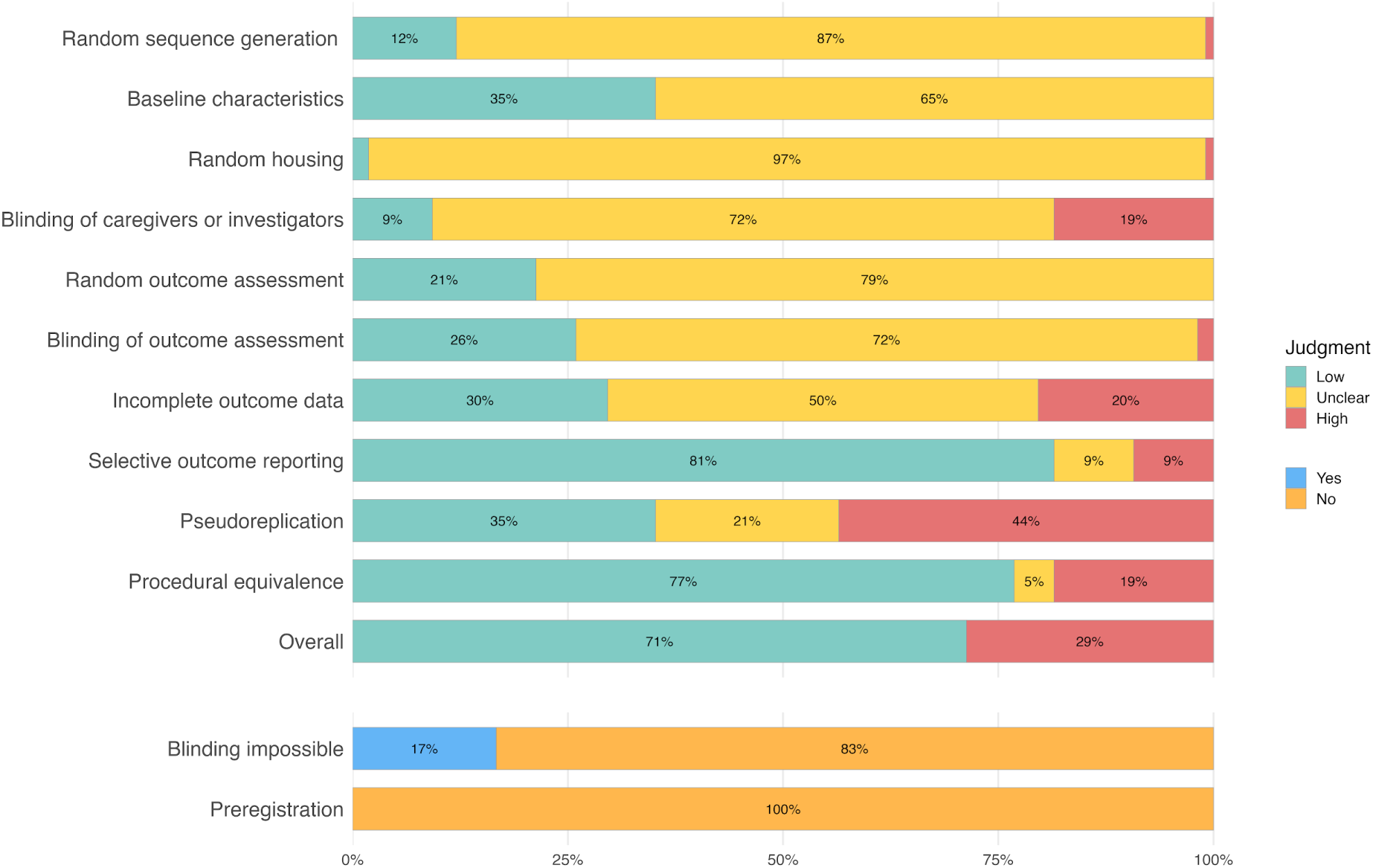
Risk of bias assessment of the included studies. Each item was scored as presenting a “low”, “unclear”, or “high” risk of bias. The possibility of blinding and study pre-registration was additionally assessed and classified as “yes” or “no”. Results are presented as the percentage of assessed studies (n = 108) presenting each score.

Overall, 71 studies had at least one item rated as high risk and 31 had two or more items rated as high risk. The risk of bias ratings assigned to each study across all assessed items are presented in Supplementary Table S3.

### 3.3. Social preference

The outcomes extracted were time in the social zone (or change from baseline), time in the non-social zone, distance from the animal to the social stimulus (or change from baseline), and distance moved within the social zone. Measures based on zone entries were not considered, as they may reflect locomotion rather than social preference.

Seven meta-analyses were performed across distinct intervention categories: anxiolytics, pro-dopaminergic agents, estradiol and related compounds, ethanol, NMDA antagonists, valproic acid, and stress (Figure 3). The results indicate an overall reduction in social preference for pro-dopaminergic agents, ethanol, NMDA antagonists, and stress-related interventions, with statistically significant pooled effects. However, the corresponding prediction intervals were wide and encompassed the null value, consistent with substantial between-study heterogeneity and indicating that future studies could observe effects in either direction.

**Figure 3.**
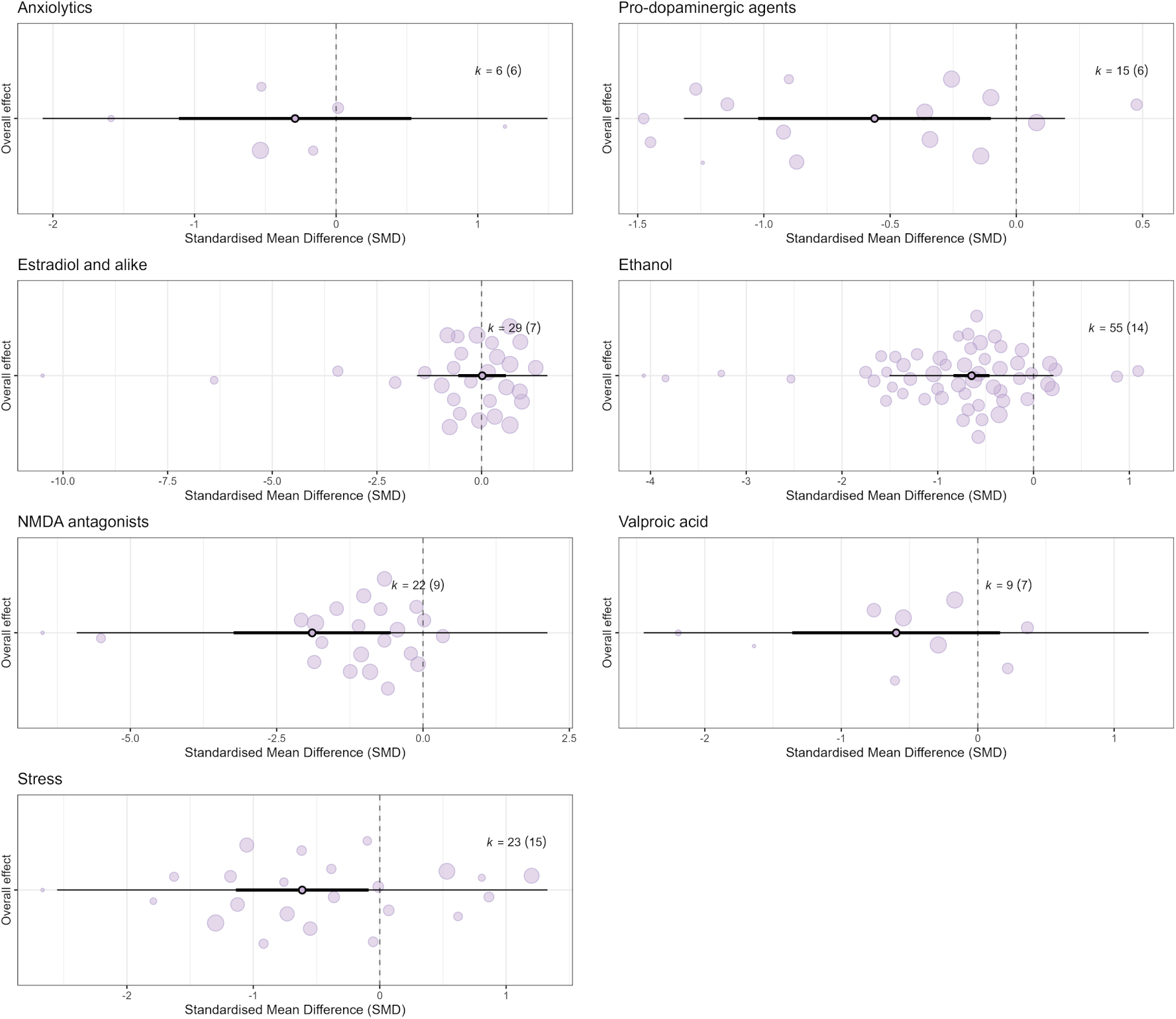
Orchard plots of social preference outcomes in adult zebrafish. The central points represent the pooled effect sizes, with thick horizontal lines indicating 95% confidence intervals and thin horizontal lines representing prediction intervals. The size of each bubble corresponds to the precision of individual effect sizes. The number of effect sizes (*k*) and the number of independent studies are shown for each category: anxiolytics, pro-dopaminergic agents, estradiol and alike, ethanol, NMDA antagonists, valproic acid, and stress. Positive values (right side) represent an increase in social preference, whereas negative values (left side) represent a decrease in social preference.

#### 3.3.1. Anxiolytics

The meta-analysis comprised 6 comparisons out of 6 independent studies and showed no significant effects on social preference (SMD: -0.29; CI 95%: -1.11, 0.53; prediction interval: -2.07, 1.49).

Substantial heterogeneity was observed (Q = 12.53, df = 5, p = 0.0282; I² = 63.57%), with 31.79% of the total variance attributable to both between- and within-study variation (σ²_between_ = 0.1896, σ²_within_ = 0.1896).

No evidence of small-study effects was detected based on visual inspection of the funnel plot (Supplementary Figure S1) or the Egger-type regression based on standard error (Table 1), also illustrated as a bubble plot (Figure 4).

**Figure 4.**
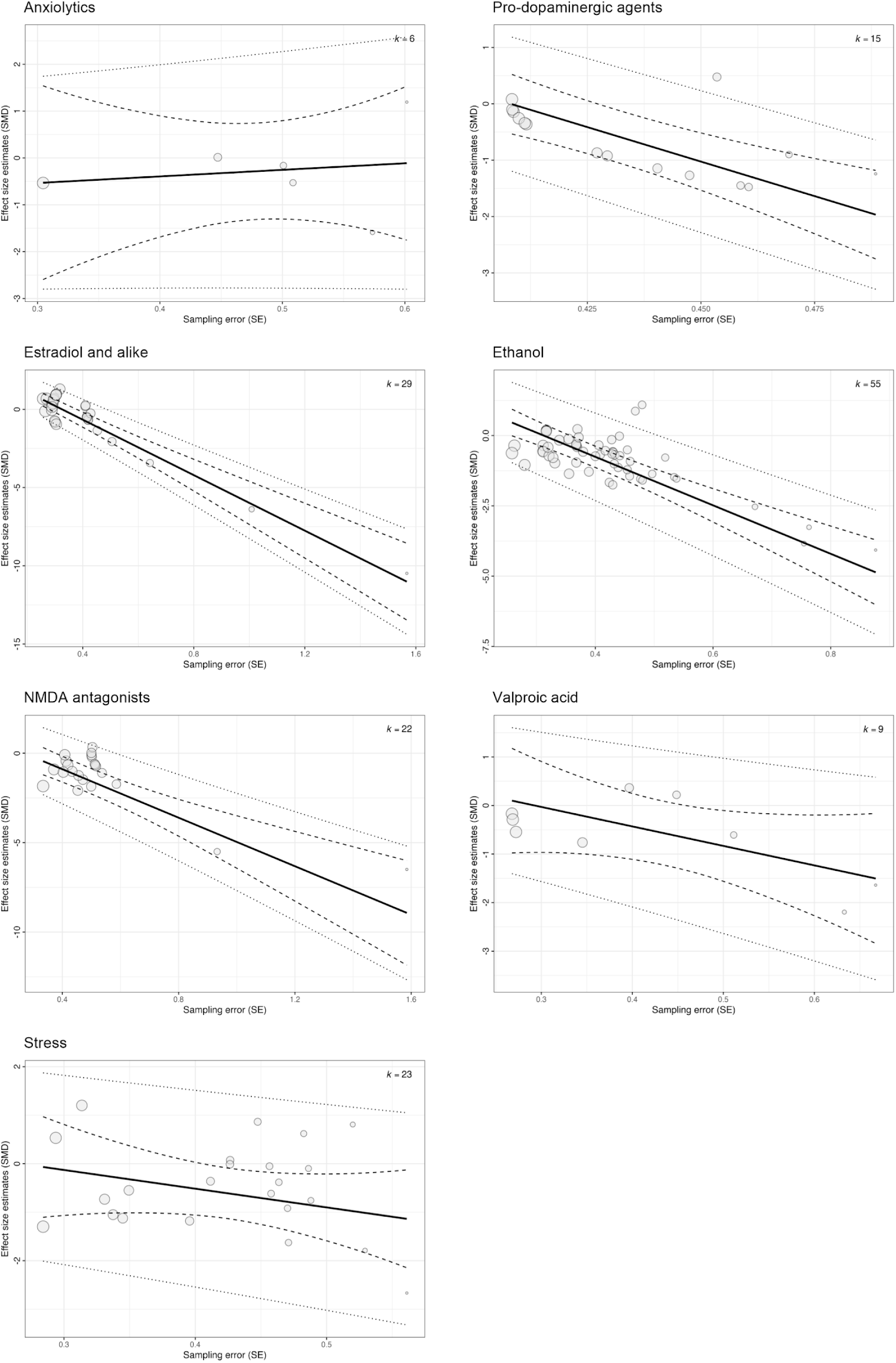
Bubble plot of social preference outcomes in adult zebrafish. The plot illustrates the relationship between sampling error (SE) and the estimated effect size (SMD). The solid black line represents the predicted average effect across the range of study precision, while the dashed and dotted lines indicate the 95% confidence and prediction intervals, respectively. Each bubble represents an individual effect size, with the size of the bubble proportional to its weight in the meta-analytical model. A visually non-zero slope is consistent with Egger regression evidence for small-study effects; a near-zero slope supports their absence.

**Table 1.**
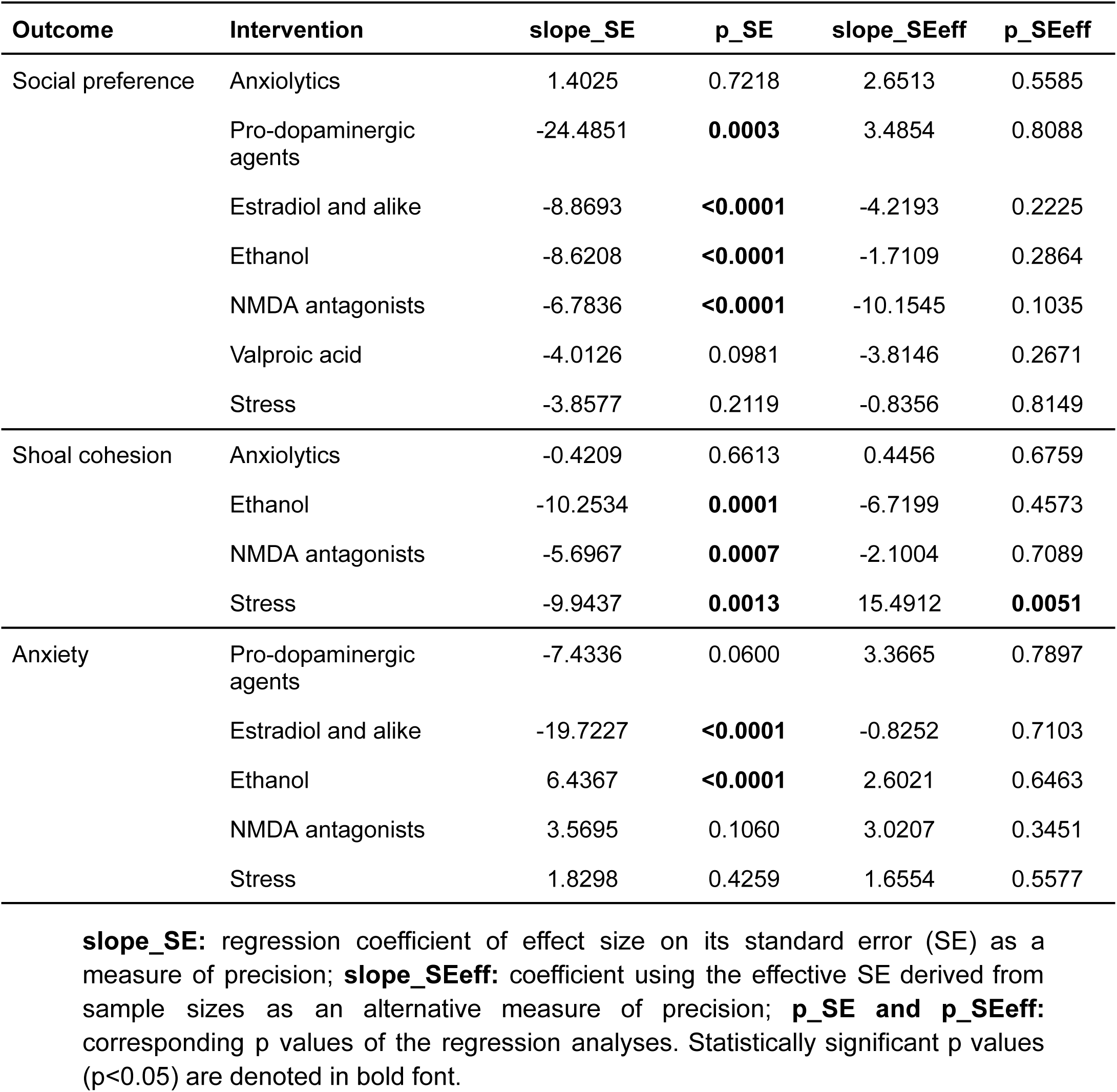
Egger-type regression analyses to assess small-study effects.

Leave-one-study-out analyses showed that the overall estimate remained non-significant after the removal of each study, except for Barba-Escobedo et al. (2012), whose exclusion yielded a statistically significant negative effect (SMD: -0.50; CI 95%: -0.87, -0.12); it was the only study reporting an increase in social preference. Excluding studies with a high overall risk of bias (one study) did not alter the results (SMD: −0.11; 95% CI: −0.63, 0.41). Pooled estimates were virtually unchanged across different levels of within-study correlation, indicating robustness to assumptions of within-study dependence.

#### 3.3.2. Pro-dopaminergic agents

The meta-analysis comprised 15 comparisons out of 6 independent studies and showed a significant decrease on social preference (SMD: -0.56; CI 95%: -1.02, -0.10; prediction interval -1.32, 0.19). However, this finding should be interpreted with caution due to the limited number of independent studies.

Heterogeneity was observed (Q = 37.42, df = 14, p = 0.0006; I² = 22.36%), with 11.18% of the total variance attributable to both between- and within-study variation (σ²_between_ = 0.0271, σ²_within_ = 0.0271).

Evidence of small-study effects was detected based on visual inspection of the funnel plot (Supplementary Figure S1) and the Egger-type regression based on standard error (Table 1), also illustrated as a bubble plot (Figure 4). However, this should be interpreted with caution as the Egger-type regression based on sample size did not confirm this finding.

Leave-one-study-out analyses showed that the overall estimate remained significant after the removal of each study. Excluding studies with a high overall risk of bias (one study) did not alter the results (SMD: −0.66; 95% CI: −1.00, -0.33). Pooled estimates were virtually unchanged across different levels of within-study correlation, indicating robustness to assumptions of within-study dependence.

#### 3.3.3. Estradiol and alike

The meta-analysis comprised 29 comparisons out of 7 independent studies and showed no significant effects on social preference (SMD: 0.02; CI 95%: -0.55, 0.59, prediction interval -1.54, 1.58).

Heterogeneity was observed (Q = 200.44, df = 28, p < 0.0001; I^2^ = 74.94%), with 26.64% of the variance attributable to between-study and 48.30% attributable to within-study variation (σ²_between_ = 0.1254, σ²_within_ = 0.2274).

Evidence of small-study effects was detected based on visual inspection of the funnel plot (Supplementary Figure S1) and the Egger-type regression based on standard error (Table 1), also illustrated as a bubble plot (Figure 4). However, this should be interpreted with caution as the Egger-type regression based on sample size did not confirm this finding.

Leave-one-study-out analyses showed that the overall estimate remained non-significant after the removal of each study. Excluding studies with a high overall risk of bias (three studies) did not alter the results (SMD: −0.20; 95% CI: −1.00, 0.60). Pooled estimates were virtually unchanged across different levels of within-study correlation, indicating robustness to assumptions of within-study dependence.

#### 3.3.4. Ethanol

The meta-analysis comprised 55 comparisons out of 14 independent studies and showed a significant decrease on social preference (SMD: -0.65; CI 95%: -0.84, -0.46; prediction interval -1.50, 0.21).

Heterogeneity was observed (Q = 166.26, df = 54, p < 0.0001, I^2^ = 49.67%), mostly attributable to within-study variation (σ²_between_ < 0.0001, σ²_within_ = 0.1493).

Evidence of small-study effects was detected based on visual inspection of the funnel plot (Supplementary Figure S1) and the Egger-type regression based on standard error (Table 1), also illustrated as a bubble plot (Figure 4). However, this should be interpreted with caution as the Egger-type regression based on sample size did not confirm this finding.

Leave-one-study-out analyses showed that the overall estimate remained significant after the removal of each study. Excluding studies with a high overall risk of bias (four studies) did not alter the results (SMD: −0.64; 95% CI: −0.91, -0.36). Pooled estimates were virtually unchanged across different levels of within-study correlation, indicating robustness to assumptions of within-study dependence.

A multilevel meta-regression was conducted to examine whether the magnitude of the social preference effect varied as a function of ethanol concentration administered in zebrafish (Figure 5). Ethanol concentration significantly moderated the effect size (F(1,12) = 9.17, p = 0.0105), with higher concentrations associated with greater reductions in social preference (β = −0.80; 95% CI: −1.37, −0.22). However, heterogeneity remained significant (Q = 136.25, df = 53, p < 0.0001).

**Figure 5.**
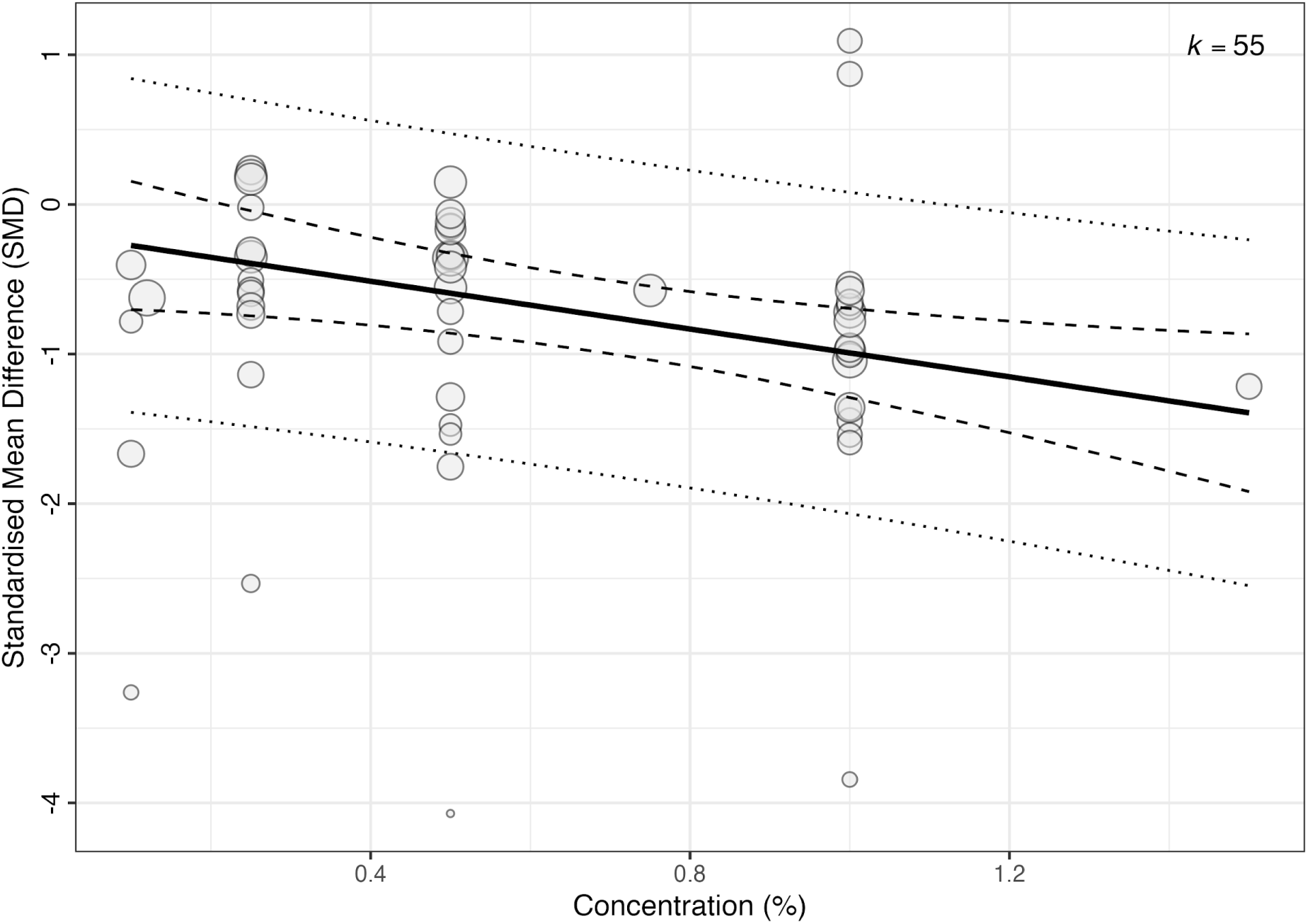
Meta-regression of ethanol effects on social preference with ethanol concentration as a moderator. Each point represents an effect size, with clustering by study. For each one–percentage point increase in ethanol concentration, the SMD decreases by 0.80 units, consistent with a dose-dependent reduction in social preference.

#### 3.3.5. NMDA antagonists

The meta-analysis comprised 22 comparisons out of 9 independent studies and showed a significant decrease on social preference (SMD: -1.89; CI 95%: -3.24, -0.55; prediction interval -5.92, 2.13).

Heterogeneity was observed (Q = 83.44, df = 21, p < 0.0001; I^2^ = 92.33%), mostly attributable to between-study variation (σ²_between_ = 2.7046, σ²_within_ < 0.0001).

Evidence of small-study effects was detected based on visual inspection of the funnel plot (Supplementary Figure S1) and the Egger-type regression based on standard error (Table 1), also illustrated as a bubble plot (Figure 4). However, this should be interpreted with caution as the Egger-type regression based on sample size did not confirm this finding.

Leave-one-study-out analyses showed that the overall estimate remained significant after the removal of each study. Excluding studies with a high overall risk of bias (one study) did not alter the results (SMD: −2.11; 95% CI: −3.39, -0.82). Pooled estimates were virtually unchanged across different levels of within-study correlation, indicating robustness to assumptions of within-study dependence.

### 3.3.6. Valproic acid

The meta-analysis comprised 9 comparisons out of 7 independent studies and showed no significant effects on social preference (SMD: -0.60; CI 95%: -1.36, 0.16; prediction interval -2.45, 1.25).

Heterogeneity was observed (Q = 20.44, df = 8, p = 0.0088; I^2^ = 77.81%), with 38.90% of the variance attributable to both between- and within-study variation (σ²_between_ = 0.2373, σ²_within_ = 0.2373).

No evidence of small-study effects was detected based on visual inspection of the funnel plot (Supplementary Figure S1) or the Egger-type regression based on standard error (Table 1), also illustrated as a bubble plot (Figure 4).

Leave-one-study-out analyses showed that the overall estimate was affected by the removal of Chele et al. (2024) and Liu et al. (2021), whose exclusion yielded a statistically significant negative effect (SMD: -0.74 CI 95%: -1.41, -0.07) and (SMD: -0.76 CI 95%: -1.38, -0.14), respectively. No study presented a high overall risk of bias within this category to be excluded. Pooled estimates were virtually unchanged across different levels of within-study correlation, indicating robustness to assumptions of within-study dependence.

#### 3.3.7. Stress

The meta-analysis comprised 23 comparisons out of 15 independent studies and showed a significant decrease on social preference (SMD: -0.61; CI 95%: -1.14, -0.09; prediction interval -2.55, 1.33).

Heterogeneity was observed (Q = 108.20, df = 22, p < 0.0001; I^2^ = 82.43%), with 67.68% of the variance attributable to between-study and 14.75% attributable to within-study variation (σ²_between_ = 0.6227, σ²_within_ = 0.1357).

No evidence of small-study effects was detected based on visual inspection of the funnel plot (Supplementary Figure S1) or the Egger-type regression based on standard error (Table 1), also illustrated as a bubble plot (Figure 4).

Leave-one-study-out analyses showed that the overall estimate remained significant after the removal of each study. Excluding studies with a high overall risk of bias (12 studies) changed the pooled estimate from a significant decrease to a non-significant effect (SMD: −0.02; 95% CI: −1.04, 1.00) based, however, on a limited number of studies (n = 3). Pooled estimates were virtually unchanged across different levels of within-study correlation, indicating robustness to assumptions of within-study dependence.

### 3.4. Shoal cohesion

The outcomes extracted were interfish distance between shoal members, nearest neighbor distance, shoal area, dispersion index, and latency to shoal formation. Four meta-analyses were conducted across distinct intervention categories: anxiolytics, ethanol, NMDA antagonists, and stress (Figure 6). Stress-related interventions were associated with an increase in shoal cohesion; however, the prediction interval crossed the null value.

**Figure 6.**
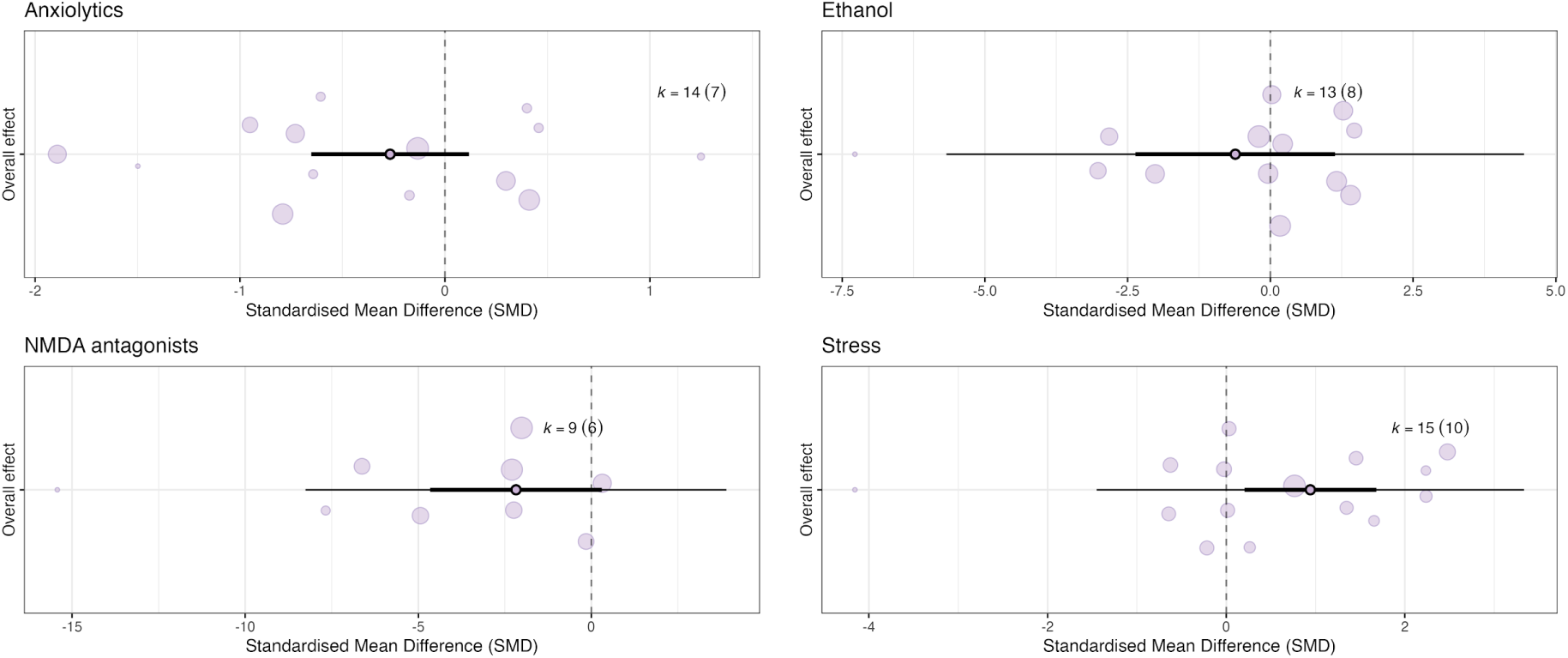
Orchard plots of shoal cohesion outcomes in adult zebrafish. The central points represent the pooled effect sizes, with thick horizontal lines indicating 95% confidence intervals and thin horizontal lines representing prediction intervals. The size of each bubble corresponds to the precision of individual effect sizes. The number of effect sizes (*k*) and the number of independent studies are shown for each category: anxiolytics, ethanol, NMDA antagonists and stress. Positive values (right side) represent an increase in shoal cohesion, whereas negative values (left side) represent a decrease in shoal cohesion.

#### 3.4.1. Anxiolytics

The meta-analysis comprised 14 comparisons out of 7 independent studies and showed no significant effects on shoal cohesion (SMD: -0.27; CI 95%: -0.65, 0.12; prediction interval -0.65, 0.12).

Heterogeneity was observed (Q = 39.24, df = 13, p = 0.0002; I² < 0.0001). However, variance component estimates indicated that no residual heterogeneity was attributable to either between-study or within-study variation, with both variance parameters estimated as zero (σ²_between_ = 0, σ²_within_ = 0).

No evidence of small-study effects was detected based on visual inspection of the funnel plot (Supplementary Figure S2) or the Egger-type regression based on standard error (Table 1), also illustrated as a bubble plot (Figure 7).

**Figure 7.**
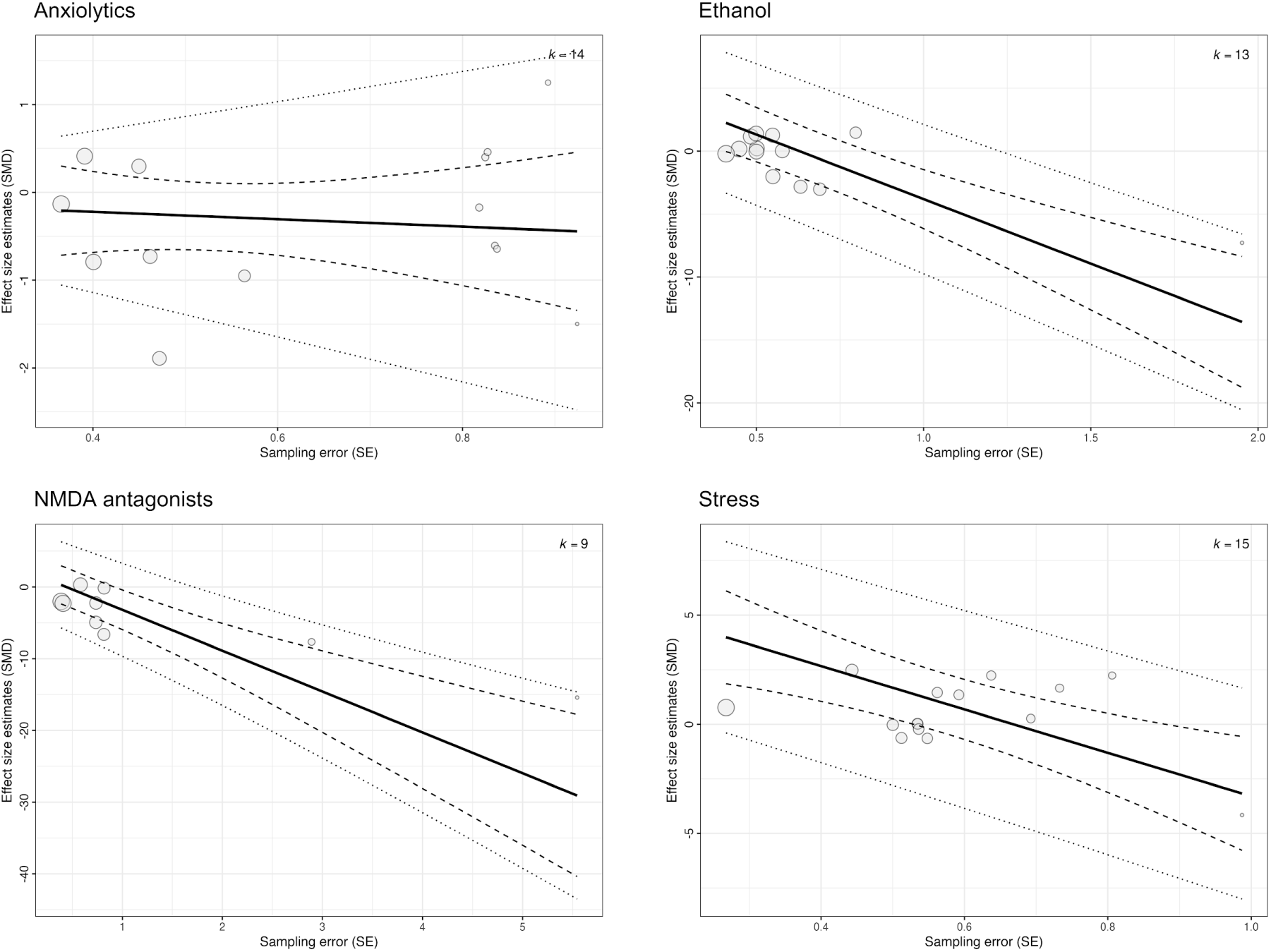
Bubble plot of shoal cohesion outcomes in adult zebrafish. The plot illustrates the relationship between sampling error (SEM) and the estimated effect size (SMD). The solid black line represents the predicted average effect across the range of study precision, while the dashed and dotted lines indicate the 95% confidence and prediction intervals, respectively. Each bubble represents an individual effect size, with the size of the bubble proportional to its weight in the meta-analytical model. A visually non-zero slope is consistent with Egger regression evidence for small-study effects; a near-zero slope supports their absence.

Leave-one-study-out analyses showed that the overall estimate remained non-significant after the removal of each study, except for Maaswinkel et al. (2013_B), whose exclusion yielded a statistically significant negative effect (SMD: -0.42; CI 95%: -0.82, -0.03). No study presented a high overall risk of bias within this category to be excluded. Pooled estimates were virtually unchanged across different levels of within-study correlation, indicating robustness to assumptions of within-study dependence.

#### 3.4.2. Ethanol

The meta-analysis comprised 13 comparisons out of 8 independent studies and showed no significant effects on shoal cohesion (SMD: -0.62; CI 95%: -2.36, 1.13; prediction interval -5.67, 4.44).

Heterogeneity was observed (Q = 122.53, df = 12, p < 0.0001; I^2^ = 92.97%), with 46.93% of the variance attributable to between-study and 46.03% attributable to within-study variation (σ²_between_ = 2.0344, σ²_within_ = 1.9956).

Evidence of small-study effects was detected based on visual inspection of the funnel plot (Supplementary Figure S2) and the Egger-type regression based on standard error (Table 1), also illustrated as a bubble plot (Figure 7). However, this should be interpreted with caution as the Egger-type regression based on sample size did not confirm this finding.

Leave-one-study-out analyses showed that the overall estimate remained non-significant after the removal of each study. Excluding studies with a high overall risk of bias (one study) did not alter the results (SMD: −0.76; 95% CI: −2.56, 1.03). Pooled estimates were virtually unchanged across different levels of within-study correlation, indicating robustness to assumptions of within-study dependence.

#### 3.4.3. NMDA antagonists

The meta-analysis comprised 9 comparisons out of 6 independent studies and showed no significant effects on shoal cohesion (SMD: -2.18; CI 95%: -4.66, 0.31; prediction interval -8.26, 3.90).

Heterogeneity was observed (Q = 86.61, df = 8, p < 0.0001; I^2^ = 91.30%), with 45.65% of the total variance attributable to both between- and within-study variation (σ²_between_ = 2.3314, σ²_within_ = 2.3314).

Evidence of small-study effects was detected based on visual inspection of the funnel plot (Supplementary Figure S2) and the Egger-type regression based on standard error (Table 1), also illustrated as a bubble plot (Figure 7). However, this should be interpreted with caution as the Egger-type regression based on sample size did not confirm this finding.

Leave-one-study-out analyses showed that the overall estimate was affected by the removal of Mccutcheon et al. (2016) and Kyzar et al. (2016), whose exclusion yielded a statistically significant negative effect (SMD = −2.72; 95% CI: −4.73, −0.72, and SMD = −2.65; 95% CI: −4.86, −0.43, respectively). Excluding studies with a high overall risk of bias (two studies) changed the pooled estimate from non-significant to a statistically significant negative effect (SMD: −2.11; 95% CI: −4.20, -0.02). Pooled estimates were virtually unchanged across different levels of within-study correlation, indicating robustness to assumptions of within-study dependence.

#### 3.4.4. Stress

The meta-analysis comprised 15 comparisons out of 10 independent studies and showed a significant increase on shoal cohesion (SMD: 0.94; CI 95%: 0.20, 1.68; prediction interval -1.45, 3.33).

Heterogeneity was observed (Q = 73.18, df = 14, p < 0.0001; I^2^ = 78.22%), mostly attributable to within-study variation (σ²_between_ < 0.0001, σ²_within_ = 1.0124).

Evidence of small-study effects was detected based on visual inspection of the funnel plot and driven by a single study (Supplementary Figure S2), and confirmed by the Egger-type regression based on standard error (Table 1), also illustrated as a bubble plot (Figure 7). However, in the Egger-type regression based on sample size the slope direction changed after adjustment. Stress was the only intervention showing a reversal in slope when the adapted standard error based on effective sample size was applied, and the association was no longer statistically significant. Overall, this pattern suggests the signal of small-study effects may be artefactual, likely driven by the known dependency between SMD and its conventional standard error.

Leave-one-study-out analyses showed that the overall estimate remained significant after the removal of each study. Excluding studies with an high overall risk of bias (seven studies) did not alter the results (SMD: 1.87; 95% CI: 1.12, 2.61). Pooled estimates were virtually unchanged across different levels of within-study correlation, indicating robustness to assumptions of within-study dependence.

### 3.5. Anxiety-related behavior

Anxiety-related measures were extracted as a secondary outcome when available. The outcomes extracted were time in the upper zone, time in the bottom zone, average height of the animal in the tank, latency to entry the upper zone, freezing time, and entries to the upper zone for the novel tank test; time in the light zone, time in the dark zone, and latency to enter the dark zone for the light/dark test; time in the center zone, time in the peripheral zone, and freezing time for the open tank test. Five meta-analyses were conducted across distinct intervention categories: pro-dopaminergic agents, estradiol and related compounds, ethanol, NMDA antagonists, and stress (Figure 8).

**Figure 8.**
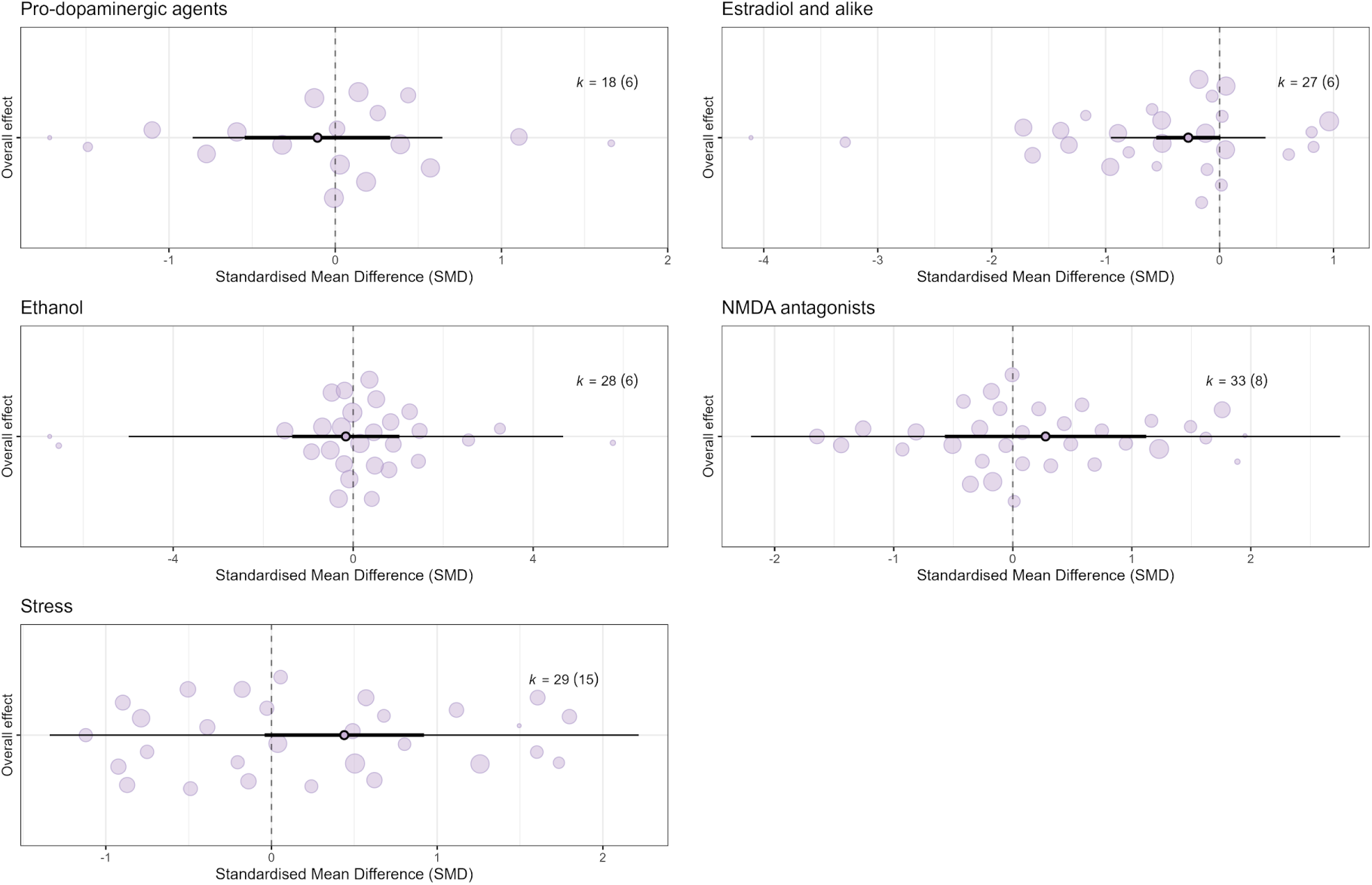
Orchard plots of anxiety related outcomes in adult zebrafish. The central points represent the pooled effect sizes, with thick horizontal lines indicating 95% confidence intervals and thin horizontal lines representing prediction intervals. The size of each bubble corresponds to the precision of individual effect sizes. The number of effect sizes (*k*) and the number of independent studies are shown for each category: pro-dopaminergic agents, estradiol and alike, ethanol, NMDA antagonists and stress. Positive values (right side) represent an increase in anxiety-like behavior, whereas negative values (left side) represent a decrease in anxiety-like behavior.

#### 3.5.1. Pro-dopaminergic agents

The meta-analysis comprised 18 comparisons out of 6 independent studies and showed no significant effects on anxiety-related behavior (SMD: -0.11; CI 95%: -0.55, 0.33; prediction interval -0.86, 0.64).

Heterogeneity was observed (Q = 79.61, df = 17, p < 0.0001; I^2^ = 22.97%), mostly attributable to between-study variation (σ²_between_ = 0.0564, σ²_within_ < 0.0001).

No evidence of small-study effects was detected based on visual inspection of the funnel plot (Supplementary Figure S3) or the Egger-type regression based on standard error (Table 1), also illustrated as a bubble plot (Figure 9).

**Figure 9.**
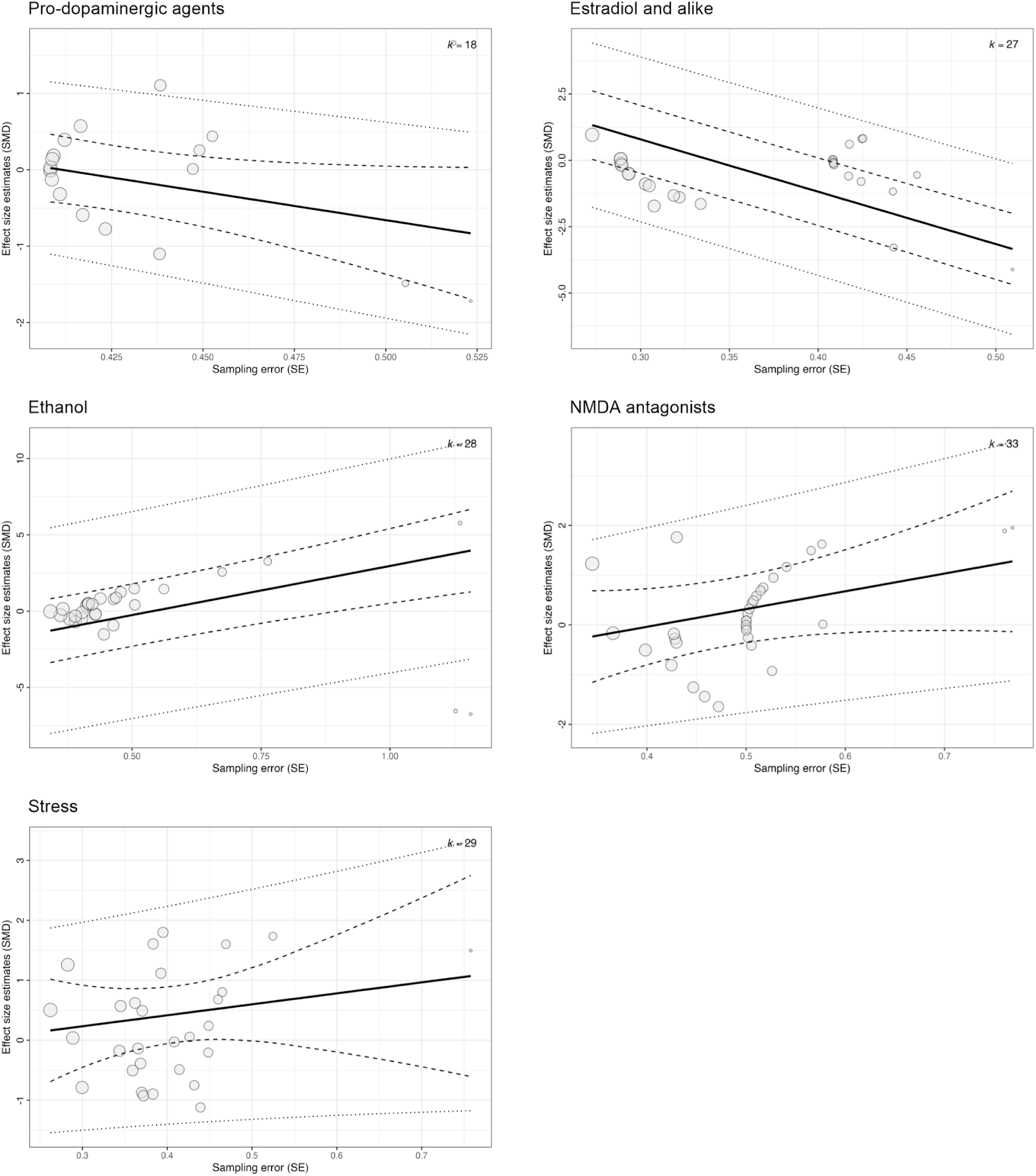
Bubble plot of anxiety related outcomes in adult zebrafish. The plot illustrates the relationship between sampling error (SEM) and the estimated effect size (SMD). The solid black line represents the predicted average effect across the range of study precision, while the dashed and dotted lines indicate the 95% confidence and prediction intervals, respectively. Each bubble represents an individual effect size, with the size of the bubble proportional to its weight in the meta-analytical model. A visually non-zero slope is consistent with Egger regression evidence for small-study effects; a near-zero slope supports their absence.

Leave-one-study-out analyses showed that the overall estimate remained non-significant after the removal of each study. Pooled estimates were virtually unchanged across different levels of within-study correlation, indicating robustness to assumptions of within-study dependence.

#### 3.5.2. Estradiol and alike

The meta-analysis comprised 27 comparisons out of 6 independent studies and showed no significant effects on anxiety-related behavior (SMD: -0.27; CI 95%: -0.56, 0.01; prediction interval -0.95, 0.40).

Heterogeneity was observed (Q = 306.57, df = 26, p < 0.0001; I^2^ = 32.05%), mostly attributable to between-study variation (σ²_between_ < 0.0001, σ²_within_ = 0.058).

Evidence of small-study effects was detected based on visual inspection of the funnel plot (Supplementary Figure S3) and the Egger-type regression based on standard error (Table 1), also illustrated as a bubble plot (Figure 9). However, this should be interpreted with caution as the Egger-type regression based on sample size did not confirm this finding.

Leave-one-study-out analyses showed that the overall estimate was affected by the removal of Fenske et al. (2020) and Reyhanian (2011), whose exclusion yielded a statistically significant negative effect (SMD = −0.37; 95% CI: −0.68, −0.06, and SMD = −0.31; 95% CI: −0.62, −0.01, respectively). The effect of estradiol and alike on anxiety was statistically significant at low ρ values (ρ = 0.0-0.4), but remained non-significant when assuming within-study dependence of ρ = 0.5-0.8.

#### 3.5.3. Ethanol

The meta-analysis comprised 28 comparisons out of 6 independent studies and showed no significant effects on anxiety-related behavior (SMD: -0.16; CI 95%: -1.36, 1.03; prediction interval -4.99, 4.66).

Heterogeneity was observed (Q = 148.61, df = 27, p < 0.0001; I^2^ = 94.11%), mostly attributable to within-study variation (σ²_between_ < 0.0001, σ²_within_ = 3.3119).

Evidence of small-study effects was detected based on visual inspection of the funnel plot (Supplementary Figure S3) and the Egger-type regression based on standard error (Table 1), also illustrated as a bubble plot (Figure 9). However, this should be interpreted with caution as the Egger-type regression based on sample size did not confirm this finding.

Leave-one-study-out analyses showed that the overall estimate remained non-significant after removal of each study. Pooled estimates were virtually unchanged across different levels of within-study correlation, indicating robustness to assumptions of within-study dependence.

#### 3.5.4. NMDA antagonists

The meta-analysis comprised 33 comparisons out of 8 independent studies and showed no significant effects on anxiety-related behavior (SMD: 0.28; CI 95%: -0.57, 1.12; prediction interval -2.20, 2.75).

Heterogeneity was observed (Q = 94.96, df = 32, p < 0.0001; I^2^ = 80.56%), with 63.77% of the variance attributable to between-study and 16.79% attributable to within-study variation (σ²_between_ = 0.7656, σ²_within_ = 0.2015).

No evidence of small-study effects was detected based on visual inspection of the funnel plot (Supplementary Figure S3) or the Egger-type regression based on standard error (Table 1), also illustrated as a bubble plot (Figure 9).

Leave-one-study-out analyses showed that the overall estimate remained non-significant after the removal of each study. Pooled estimates were virtually unchanged across different levels of within-study correlation, indicating robustness to assumptions of within-study dependence.

#### 3.5.5. Stress

The meta-analysis comprised 29 comparisons out of 15 independent studies and showed no significant effects on anxiety-related behavior (SMD: 0.44; CI 95%: -0.04, 0.92; prediction interval -1.34, 2.22).

Heterogeneity was observed (Q = 146.92, df = 28, p < 0.0001; I^2^ = 81.58%), with 72.74% of the variance attributable to between-study and 8.83% attributable to within-study variation (σ²_between_ = 0.5679, σ²_within_ = 0.0690).

No evidence of small-study effects was detected based on visual inspection of the funnel plot (Supplementary Figure S3) or the Egger-type regression based on standard error (Table 1), also illustrated as a bubble plot (Figure 9).

Leave-one-study-out analyses showed that the overall estimate was affected by the removal of Barcellos et al. (2020), Tamagno et al. (2022_A), Marchetto et al. (2021), Müller et al. (2024) and Shams et al. (2017_A), whose exclusion yielded a statistically significant positive effect. Pooled estimates were virtually unchanged across different levels of within-study correlation, indicating robustness to assumptions of within-study dependence.

## 4. Discussion

This systematic review and meta-analysis synthesized the effects of pharmacological interventions that modulate the central nervous system and stress-related interventions on adult zebrafish social behavior and, when available, anxiety-like behavior. Zebrafish that were exposed to interventions classified as pro-dopaminergic agents, ethanol, NMDA antagonists, and stress-related showed decreased social behavior in the social preference test; at the same time, stress-related interventions increased social behavior in the shoal cohesion test.

Social preference and shoal cohesion were analyzed in separate meta-analyses because these paradigms likely capture distinct components of social behavior. The social preference test is conceptually analogous to the rodent three-chamber paradigm and primarily assesses motivation for social approach toward conspecifics. In contrast, shoal cohesion reflects a species-specific collective behavior in fish, integrating social attraction with sensory processing, motor coordination, and group-level dynamics (Kalueff et al., 2014). Treating these outcomes separately, therefore, improves construct validity and reduces the risk of conflating mechanistically distinct social phenotypes. The diverging effects observed for stress-related interventions further supports these assays as complementary rather than interchangeable outcomes.

The reduction of social preference after treatment with pro-dopaminergic agents and NMDA antagonists may reflect the dysfunctional hyperdopaminergic state that compromises social behavior, as observed in psychosis-related conditions (Benvenutti et al., 2022; Dichter et al., 2012). In contrast, our analysis yielded no significant effects for such interventions on shoal cohesion or anxiety-related behaviors. While previous studies have reported that NMDA receptor antagonists reduce shoal cohesion in zebrafish (Riehl et al., 2011; Seibt et al., 2010), these compounds have also been described as exerting anxiolytic effects (De Campos et al., 2015; Maaswinkel et al., 2013a; Riehl et al., 2011). Because a decrease in anxiety levels can independently reduce shoaling behavior, shoal cohesion may be confounded when used as a measure of social withdrawal to model behavioral phenotypes relevant to neuropsychiatric disorders. Since the current data did not indicate robust changes in anxiety-like behavior, it remains unclear whether the previously reported disruptions in shoal cohesion are secondary to anxiolysis or if they represent distinct, socially driven mechanisms. Further experimental studies are thus warranted to disentangle the potential influence of anxiolytic and anxiogenic interventions on shoal cohesion, and to differentiate true social alterations from secondary consequences of anxiety modulation.

Our meta-analytic findings indicate that ethanol exposure is associated with a reduction of social preference in zebrafish in a dose-dependent manner, with higher ethanol concentrations being associated with a greater reduction in social preference. No significant effect was observed for shoal cohesion, suggesting that ethanol impairs social motivation without necessarily disrupting group-level organization. The effects of ethanol on anxiety-like behavior are not well elucidated in the literature. While low to moderate doses have been shown to exert anxiolytic effects, higher doses are associated with motor impairments (Clayman & Connaughton, 2022; Kameda et al., 2007; Miranda-Morales et al., 2014). This dose-dependent pattern may explain why no significant effects on anxiety-like behavior were found; as few studies assessed locomotion, a clear evaluation of this potential dose-related confounder was precluded.

The finding that stress decreases social preference in zebrafish is consistent with observations across other species (van Boxelaere et al., 2017; Varlinskaya & Spear, 2012; Zain et al., 2019). Based on this pattern, a concomitant reduction in shoal cohesion would be expected. However, stress exposure increased proximity among group members, a response consistent with an adaptive aggregation mechanism that increases foraging efficiency and predator avoidance (Facciol & Gerlai, 2020). These findings are consistent with studies in which acute stress induced by predator cues or exposure to a novel environment transiently triggered a rapid dispersion of the shoal, followed by a marked increase in shoal cohesion, reflecting a protective grouping response (Kleinhappel et al., 2019; Miller & Gerlai, 2007). In another study, alarm substance exposure also increased shoal cohesion and elicited an anxiogenic-like profile in the novel tank test (Speedie & Gerlai, 2008). Similarly, a meta-analysis focusing on the effects of unpredictable chronic stress in zebrafish reported an overall anxiogenic profile without a significant effect on social behavior (Gallas-Lopes et al., 2023). In our meta-analysis, different types of stressors, including acute and chronic, were grouped; while the pooled effect was null, it was challenged in sensitivity analysis, further suggesting a trend towards increased anxiety, as expected.

Our results indicate that social preference and shoal cohesion are not interchangeable measures of social behavior. Interventions may differentially affect social approach towards conspecifics versus group-level spatial organization, reinforcing the need to interpret these assays as complementary but non-equivalent outcomes. Zebrafish social interactions, particularly in shoal cohesion assays, are context-dependent and influenced by multiple environmental and internal factors, including predator presence, individual characteristics (e.g., age or body size), stress levels, hunger, familiarity among conspecifics, and the sex composition of the shoal (Maaswinkel et al., 2013a; Miller & Gerlai, 2007; Swaney et al., 2025). Moreover, changes in shoal cohesion may be less sensitive or less specific to measure social behavior, as compared to social preference. When studying the impact of an intervention on shoal cohesion, it is important to determine whether eventual alterations primarily reflect social functioning or are secondary to modulation in anxiety levels.

In addition to biological and experimental factors, methodological aspects related to tracking resolution for spatial parameters represented an important source of variability across studies. Differences in the position of the camera, video resolution, tracking algorithms, and division of the zones can affect the precision with which inter-individual distances and spatial preferences are quantified. Moreover, the same behavioral recordings can be analyzed using multiple analytical pipelines and outcome definitions, potentially leading to variability in reported results across studies. Such methodological flexibility underscores the importance of transparency and standardization in tracking and analysis procedures when interpreting social behavior outcomes in zebrafish.

Despite some interventions being statistically significant in this review, prediction intervals of all categories encompassed the null value, indicating that the direction and magnitude of social effects are highly context-dependent, likely influenced by factors such as duration of intervention, behavioral protocol, concentration, and environmental conditions. The significant moderating effect of ethanol concentration provides an important proof of principle: when sufficient data are available and a biologically meaningful moderator is tested, part of the observed heterogeneity can be explained. The scarcity of similar moderator analyses for other intervention classes reflects limitations in the number of studies available in each category.

### 4.2. Limitations

An important limitation of this study concerns the categorization of interventions. Although grouping was necessary for quantitative synthesis according to their nature, mechanism of action, and/or therapeutic purpose, some categories encompassed heterogeneous interventions that may not be fully captured by a single classification. This within-category heterogeneity may have increased variability and reduced sensitivity to detect specific effects, and should therefore be considered when interpreting the pooled results. However, given the small number of studies included in each category, alternative classification schemes would be unlikely to substantially alter the pooled effect estimates.

Our analyses were restricted to the available data and searched databases, and are therefore subject to publication bias, as confirmed for some interventions. Three records could not be retrieved, highlighting the importance of disseminating results through preprint platforms and other open-access channels to mitigate barriers to data access. In addition, as an observational synthesis, the associations identified do not allow causal inferences and should be interpreted as hypothesis-generating.

Throughout the review process, we identified substantial deficiencies in the reporting of important methodological details, ranging from basic animal characteristics, such as sex and exact age, to critical data elements, including exact sample sizes per experimental group and raw data availability. Consequently, data extraction often relied on information derived from figures, which may introduce minor human errors. In several cases, sample sizes had to be inferred, either by adopting the lowest value within a reported range or by estimating the number of data points displayed in graphical representations. Similarly, when error bars were not clearly visible, conservative assumptions regarding the magnitude or placement of variability estimates were made to avoid overestimation of effects. In one case, substantial overlap of data points in the graph of interest prevented reliable data extraction, thereby preventing this study from being included in this review.

### 4.3. Future directions

Together, these limitations underscore an urgent need for enhanced transparency in zebrafish behavioral research. Data sharing in open repositories and adoption of the ARRIVE guidelines (Percie Du Sert et al., 2020) for reporting could minimize these hurdles and improve the precision of evidence synthesis. Future investigations must prioritize rigorous scientific practices, specifically concerning randomization procedures, blinding, exact sample size disclosure, and the appropriate statistical handling of non-independence. Furthermore, while pre-registration of animal studies remains rare in this field, its broader adoption would substantially mitigate selective outcome reporting and bolster overall reproducibility.

Despite current methodological shortcomings in the literature, our findings validate adult zebrafish as a valuable model species for investigating the neurobiological modulation of social behavior. While current evidence indicates that specific interventions effectively modulate sociality, the high heterogeneity and generally low methodological quality of available data temper the strength of these conclusions. For other interventions, the null effect was challenged in sensitivity analyses, which may also be influenced by the limited number of available studies and between-study heterogeneity. Ultimately, well-designed, rigorous research is essential to corroborate these findings and to expand the evidence base, particularly for interventions that currently lack sufficient data for quantitative synthesis.

## 5. Conclusion

This systematic review and meta-analysis showed that some pharmacological interventions that modulate the central nervous system alter social behavior in adult zebrafish, most commonly leading to reductions in social preference. Stress-related interventions differentially impact social preference and shoal cohesion, highlighting the complementary, non-interchangeable nature of these behavioral assays. However, the effects are highly heterogeneous and strongly influenced by methodological factors. The limited literature for several intervention categories constrains the strength of conclusions regarding the underlying effect. The current evidence base has some gaps that limit the breadth of current understanding and underscore the need for future research.

## 6. Author contributions

**Daniela V. Müller:** conceptualization, data curation, formal analysis, investigation, methodology, project administration, visualization, writing - original draft; writing - review and editing. **Matheus Gallas-Lopes:** conceptualization, data curation, formal analysis, investigation, methodology, project administration, visualization, writing - review and editing. **Mariana B. Abreu:** conceptualization, investigation, methodology, writing - review and editing. **Bruno D. Arbo:** conceptualization, investigation, methodology, writing - review and editing. **Leonardo M. Bastos:** conceptualization, investigation, methodology, writing - review and editing. **Nicole T. Fröhlich:** conceptualization, investigation, methodology, writing - review and editing. **Matheus Marcon:** conceptualization, investigation, methodology, writing - review and editing. **Izabela B. Moraes:** conceptualization, investigation, methodology, writing - review and editing. **Luiz Cesar C. P. da Silva:** conceptualization, investigation, methodology, writing - review and editing. **Gabriel R. Zurchimitten:** conceptualization, investigation, methodology, writing - review and editing. **Ana P. Herrmann:** conceptualization, data curation, formal analysis, funding acquisition, investigation, methodology, project administration, resources, supervision, visualization, writing - review and editing.

## 8. Data availability

Data and analysis scripts of this study are openly available via the Open Science Framework at https://osf.io/x95w6 (Gallas-Lopes et al., 2024) and via GitHub at https://github.com/BRISAcollab/social-zebrafish/ (BRISA, 2026).

## 9. Declaration of generative AI use

During the preparation of this work, the authors used ChatGPT (OpenAI, San Francisco, CA, USA; GPT-4o model) in order to assist with language editing and text refinement. After using this tool, the authors reviewed and edited the content as needed and take full responsibility for the content of the published article.

## 11. Acknowledgements

We thank Dr. Dirce Maria Santin, Librarian at Instituto de Ciências Básicas da Saúde (UFRGS), for her help reviewing the search strategies. We also thank the BRISA collaboration for their continuous support and valuable feedback throughout all stages of the project.

## References

Apukhtin, K. V., Riga, V. D., Kotova, M. M., Galstyan, D. S., Yang, L., Cui, J., Lim, L. W., S de Abreu, M., & Kalueff, A. V. (2026). Neurobiology and neuropharmacology of zebrafish social behavior. Journal of Psychopharmacology, 2698811251413496. 10.1177/02698811251413496

Avey, M. T., Moher, D., Sullivan, K. J., Fergusson, D., Griffin, G., Grimshaw, J. M., Hutton, B., Lalu, M. M., Macleod, M., Marshall, J., Mei, S. H. J., Rudnicki, M., Stewart, D. J., Turgeon, A. F., McIntyre, L., & Canadian Critical Care Translational Biology Group. (2016). The Devil Is in the Details: Incomplete Reporting in Preclinical Animal Research. PloS One, 11(11), e0166733. 10.1371/journal.pone.0166733

Benvenutti, R., Gallas-Lopes, M., Marcon, M., Reschke, C. R., Herrmann, A. P., & Piato, A. (2022). Glutamate NMDA Receptor Antagonists with Relevance to Schizophrenia:A Review of Zebrafish Behavioral Studies. Current Neuropharmacology, 20(3), 494–509. 10.2174/1570159X19666210215121428

BRISA. (2026). BRISAcollab/social-zebrafish: V1.0.0 [Software]. Zenodo. 10.5281/zenodo.19369353

Chen, P., & Hong, W. (2018). Neural Circuit Mechanisms of Social Behavior. Neuron, 98(1), 16–30. 10.1016/j.neuron.2018.02.026

Clayman, C. L., & Connaughton, V. P. (2022). Neurochemical and Behavioral Consequences of Ethanol and/or Caffeine Exposure: Effects in Zebrafish and Rodents. Current Neuropharmacology, 20(3), 560–578. 10.2174/1570159X19666211111142027

De Campos, E. G., Bruni, A. T., & De Martinis, B. S. (2015). Ketamine induces anxiolytic effects in adult zebrafish: A multivariate statistics approach. Behavioural Brain Research, 292, 537–546. 10.1016/j.bbr.2015.07.017

de Vries, R. B. M., Hooijmans, C. R., Langendam, M. W., van Luijk, J., Leenaars, M., Ritskes-Hoitinga, M., & Wever, K. E. (2015). A protocol format for the preparation, registration and publication of systematic reviews of animal intervention studies. Evidence-Based Preclinical Medicine, 2(1), e00007. 10.1002/ebm2.7

Dichter, G. S., Damiano, C. A., & Allen, J. A. (2012). Reward circuitry dysfunction in psychiatric and neurodevelopmental disorders and genetic syndromes: Animal models and clinical findings. Journal of Neurodevelopmental Disorders, 4(1), 19. 10.1186/1866-1955-4-19

Facciol, A., & Gerlai, R. (2020). Zebrafish Shoaling, Its Behavioral and Neurobiological Mechanisms, and Its Alteration by Embryonic Alcohol Exposure: A Review. Frontiers in Behavioral Neuroscience, 14, 572175. 10.3389/fnbeh.2020.572175

Fontana, B. D., Müller, T. E., Cleal, M., De Abreu, M. S., Norton, W. H. J., Demin, K. A., Amstislavskaya, T. G., Petersen, E. V., Kalueff, A. V., Parker, M. O., & Rosemberg, D. B. (2022). Using zebrafish (Danio rerio) models to understand the critical role of social interactions in mental health and wellbeing. Progress in Neurobiology, 208, 101993. 10.1016/j.pneurobio.2021.101993

Gallas-Lopes, M., Bastos, L. M., Benvenutti, R., Panzenhagen, A. C., Piato, A., & Herrmann, A. P. (2023). Systematic review and meta-analysis of 10 years of unpredictable chronic stress in zebrafish. Lab Animal, 52(10), 229–246. 10.1038/s41684-023-01239-5

Gallas-Lopes, M., Müller, D., Abreu, M., Arbo, B., Bastos, L., Fröhlich, N., Marcon, M., Moraes, I., da Silva, L., & da Rocha Zurchimitten, G. (2025). Modulation of social behavior in adult zebrafish (Danio rerio): A systematic review and meta-analysis. https://osf.io/x95w6

Gallas-Lopes, M., Müller, D. V., Abreu, M., Arbo, B. D., Bastos, L. M., Fröhlich, N. T., Marcon, M., Moraes, I. B., Silva, L. C. C. P. da, & Zurchimitten, G. da R. (2024). Modulation of social behavior in adult zebrafish (Danio rerio): A systematic review and meta-analysis. https://osf.io/h4v3k/

Gebauer, D. L., Pagnussat, N., Piato, Â. L., Schaefer, I. C., Bonan, C. D., & Lara, D. R. (2011). Effects of anxiolytics in zebrafish: Similarities and differences between benzodiazepines, buspirone and ethanol. Pharmacology Biochemistry and Behavior, 99(3), 480–486. 10.1016/j.pbb.2011.04.021

Geng, Y., & Peterson, R. T. (2019). The zebrafish subcortical social brain as a model for studying social behavior disorders. Disease Models & Mechanisms, 12(8), dmm039446. 10.1242/dmm.039446

Giacomini, A. C. V. V., Abreu, M. S., Giacomini, L. V., Siebel, A. M., Zimerman, F. F., Rambo, C. L., Mocelin, R., Bonan, C. D., Piato, A. L., & Barcellos, L. J. G. (2016). Fluoxetine and diazepam acutely modulate stress induced-behavior. Behavioural Brain Research, 296, 301–310. 10.1016/j.bbr.2015.09.027

Goodson, J. L. (2005). The vertebrate social behavior network: Evolutionary themes and variations. Hormones and Behavior, 48(1), 11–22. 10.1016/j.yhbeh.2005.02.003

Gosselin, R.-D. (2021). Insufficient transparency of statistical reporting in preclinical research: A scoping review. Scientific Reports, 11(1), 3335. 10.1038/s41598-021-83006-5

Henry, J. D., von Hippel, W., Molenberghs, P., Lee, T., & Sachdev, P. S. (2016). Clinical assessment of social cognitive function in neurological disorders. Nature Reviews. Neurology, 12(1), 28–39. 10.1038/nrneurol.2015.229

Hofmann, H. A., Beery, A. K., Blumstein, D. T., Couzin, I. D., Earley, R. L., Hayes, L. D., Hurd, P. L., Lacey, E. A., Phelps, S. M., Solomon, N. G., Taborsky, M., Young, L. J., & Rubenstein, D. R. (2014). An evolutionary framework for studying mechanisms of social behavior. Trends in Ecology & Evolution, 29(10), 581–589. 10.1016/j.tree.2014.07.008

Hooijmans, C. R., Rovers, M. M., de Vries, R. B., Leenaars, M., Ritskes-Hoitinga, M., & Langendam, M. W. (2014). SYRCLE’s risk of bias tool for animal studies. BMC Medical Research Methodology, 14, 43. 10.1186/1471-2288-14-43

Kalueff, A., Stewart, A. M., & Gerlai, R. (2014). Zebrafish as an emerging model for studying complex brain disorders. Trends in Pharmacological Sciences, 35(2), 63–75. 10.1016/j.tips.2013.12.002

Kameda, S. R., Frussa-Filho, R., Carvalho, R. C., Takatsu-Coleman, A. L., Ricardo, V. P., Patti, C. L., Calzavara, M. B., Lopez, G. B., Araujo, N. P., Abílio, V. C., Ribeiro, R. de A., D’Almeida, V., & Silva, R. H. (2007). Dissociation of the effects of ethanol on memory, anxiety, and motor behavior in mice tested in the plus-maze discriminative avoidance task. Psychopharmacology, 192(1), 39–48. 10.1007/s00213-006-0684-9

Kleinhappel, T. K., Pike, T. W., & Burman, O. H. P. (2019). Stress-induced changes in group behaviour. Scientific Reports, 9(1), 17200. 10.1038/s41598-019-53661-w

Lee, Y., Kim, D., Kim, Y.-H., Lee, H., & Lee, C.-J. (2010). Improvement of pentylenetetrazol-induced learning deficits by valproic acid in the adult zebrafish. European Journal of Pharmacology, 643(2–3), 225–231. 10.1016/j.ejphar.2010.06.041

Lieberman, J. A., Perkins, D., Belger, A., Chakos, M., Jarskog, F., Boteva, K., & Gilmore, J. (2001). The early stages of schizophrenia: Speculations on pathogenesis, pathophysiology, and therapeutic approaches. Biological Psychiatry, 50(11), 884–897. 10.1016/S0006-3223(01)01303-8

Lieschke, G. J., & Currie, P. D. (2007). Animal models of human disease: Zebrafish swim into view. Nature Reviews Genetics, 8(5), Artigo 5. 10.1038/nrg2091

Maaswinkel, H., Zhu, L., & Weng, W. (2013a). Assessing Social Engagement in Heterogeneous Groups of Zebrafish: A New Paradigm for Autism-Like Behavioral Responses. PLoS ONE, 8(10), e75955. 10.1371/journal.pone.0075955

Maaswinkel, H., Zhu, L., & Weng, W. (2013b). Using an automated 3D-tracking system to record individual and shoals of adult zebrafish. Journal of Visualized Experiments: JoVE, (82), 50681. 10.3791/50681

Marotta, R., Risoleo, M. C., Messina, G., Parisi, L., Carotenuto, M., Vetri, L., & Roccella, M. (2020). The Neurochemistry of Autism. Brain Sciences, 10(3), 163. 10.3390/brainsci10030163

Miller, N., & Gerlai, R. (2007). Quantification of shoaling behaviour in zebrafish (Danio rerio). Behavioural Brain Research, 184(2), 157–166. 10.1016/j.bbr.2007.07.007

Miller, N., Greene, K., Dydinski, A., & Gerlai, R. (2013). Effects of nicotine and alcohol on zebrafish (Danio rerio) shoaling. Behavioural Brain Research, 240, 192–196. 10.1016/j.bbr.2012.11.033

Miranda-Morales, R. S., Nizhnikov, M. E., Waters, D. H., & Spear, N. E. (2014). New evidence of ethanol’s anxiolytic properties in the infant rat. Alcohol (Fayetteville, N.Y.), 48(4), 367–374. 10.1016/j.alcohol.2014.01.007

Nakagawa, S., Lagisz, M., Jennions, M. D., Koricheva, J., Noble, D. W. A., Parker, T. H., Sánchez-Tójar, A., Yang, Y., & O’Dea, R. E. (2022). Methods for testing publication bias in ecological and evolutionary meta-analyses. Methods in Ecology and Evolution, 13(1), 4–21. 10.1111/2041-210X.13724

Nakagawa, S., Lagisz, M., O’Dea, R. E., Pottier, P., Rutkowska, J., Senior, A. M., Yang, Y., & Noble, D. W. A. (2023). orchaRd 2.0: An R package for visualising meta-analyses with orchard plots. Methods in Ecology and Evolution, 14(8), 2003–2010. 10.1111/2041-210X.14152

Newman, S. W. (1999). The medial extended amygdala in male reproductive behavior. A node in the mammalian social behavior network. Annals of the New York Academy of Sciences, 877, 242–257. 10.1111/j.1749-6632.1999.tb09271.x

Norton, W. H. J. (2013). Toward developmental models of psychiatric disorders in zebrafish. Frontiers in Neural Circuits, 7, 79. 10.3389/fncir.2013.00079

Ouzzani, M., Hammady, H., Fedorowicz, Z., & Elmagarmid, A. (2016). Rayyan-a web and mobile app for systematic reviews. Systematic Reviews, 5(1), 210. 10.1186/s13643-016-0384-4

Page, M. J., McKenzie, J. E., Bossuyt, P. M., Boutron, I., Hoffmann, T. C., Mulrow, C. D., Shamseer, L., Tetzlaff, J. M., Akl, E. A., Brennan, S. E., Chou, R., Glanville, J., Grimshaw, J. M., Hróbjartsson, A., Lalu, M. M., Li, T., Loder, E. W., Mayo-Wilson, E., McDonald, S., … Moher, D. (2021). The PRISMA 2020 statement: An updated guideline for reporting systematic reviews. Revista Espanola De Cardiologia (English Ed.), 74(9), 790–799. 10.1016/j.rec.2021.07.010

Percie Du Sert, N., Hurst, V., Ahluwalia, A., Alam, S., Avey, M. T., Baker, M., Browne, W. J., Clark, A., Cuthill, I. C., Dirnagl, U., Emerson, M., Garner, P., Holgate, S. T., Howells, D. W., Karp, N. A., Lazic, S. E., Lidster, K., MacCallum, C. J., Macleod, M., … Würbel, H. (2020). The ARRIVE guidelines 2.0: Updated guidelines for reporting animal research. PLOS Biology, 18(7), e3000410. 10.1371/journal.pbio.3000410

Pustejovsky, J. E., Pekofsky, S., & Zhang, J. (2025). *clubSandwich: Cluster-Robust (Sandwich) Variance Estimators with Small-Sample Corrections* (Versão 0.6.1) [Software].https://cran.r-project.org/web/packages/clubSandwich/index.html

Qin, M., Wong, A., Seguin, D., & Gerlai, R. (2014). Induction of Social Behavior in Zebrafish: Live Versus Computer Animated Fish as Stimuli. Zebrafish, 11(3), 185–197. 10.1089/zeb.2013.0969

Riehl, R., Kyzar, E., Allain, A., Green, J., Hook, M., Monnig, L., Rhymes, K., Roth, A., Pham, M., Razavi, R., DiLeo, J., Gaikwad, S., Hart, P., & Kalueff, A. V. (2011). Behavioral and physiological effects of acute ketamine exposure in adult zebrafish. Neurotoxicology and Teratology, 33(6), 658–667. 10.1016/j.ntt.2011.05.011

Robinson, K. J., Bosch, O. J., Levkowitz, G., Busch, K. E., Jarman, A. P., & Ludwig, M. (2019). Social creatures: Model animal systems for studying the neuroendocrine mechanisms of social behaviour. Journal of Neuroendocrinology, 31(12), e12807. 10.1111/jne.12807

Ruhl, N., McRobert, S. P., & Currie, W. J. S. (2009). Shoaling preferences and the effects of sex ratio on spawning and aggression in small laboratory populations of zebrafish (Danio rerio). Lab Animal, 38(8), 264–269. 10.1038/laban0809-264

Saverino, C., & Gerlai, R. (2008). The social zebrafish: Behavioral responses to conspecific, heterospecific, and computer animated fish. Behavioural brain research, 191(1), 77–87. 10.1016/j.bbr.2008.03.013

Savio, L. E. B., Vuaden, F. C., Piato, A. L., Bonan, C. D., & Wyse, A. T. S. (2012). Behavioral changes induced by long-term proline exposure are reversed by antipsychotics in zebrafish. Progress in Neuro-Psychopharmacology & Biological Psychiatry, 36(2), 258–263. 10.1016/j.pnpbp.2011.10.002

Schaefer, I. C., Siebel, A. M., Piato, A. L., Bonan, C. D., Vianna, M. R., & Lara, D. R. (2015). The side-by-side exploratory test: A simple automated protocol for the evaluation of adult zebrafish behavior simultaneously with social interaction. Behavioural Pharmacology, 26(7 Spec No), 691–696. 10.1097/FBP.0000000000000145

Seibt, K. J., Oliveira, R. da L., Zimmermann, F. F., Capiotti, K. M., Bogo, M. R., Ghisleni, G., & Bonan, C. D. (2010). Antipsychotic drugs prevent the motor hyperactivity induced by psychotomimetic MK-801 in zebrafish (Danio rerio). Behavioural Brain Research, 214(2), 417–422. 10.1016/j.bbr.2010.06.014

Sison, M., Cawker, J., Buske, C., & Gerlai, R. (2006). Fishing for genes influencing vertebrate behavior: Zebrafish making headway. Lab Animal, 35(5), 33–39. 10.1038/laban0506-33

Soares, M. C., Gerlai, R., & Maximino, C. (2018). The integration of sociality, monoamines and stress neuroendocrinology in fish models: Applications in the neurosciences. Journal of Fish Biology, 93(2), 170–191. 10.1111/jfb.13757

Speedie, N., & Gerlai, R. (2008). Alarm substance induced behavioral responses in zebrafish (Danio rerio). Behavioural Brain Research, 188(1), 168–177. 10.1016/j.bbr.2007.10.031

Spence, R., Gerlach, G., Lawrence, C., & Smith, C. (2008). The behaviour and ecology of the zebrafish, Danio rerio. Biological Reviews, 83(1), 13–34. 10.1111/j.1469-185X.2007.00030.x

Stewart, A. M., Braubach, O., Spitsbergen, J., Gerlai, R., & Kalueff, A. V. (2014). Zebrafish models for translational neuroscience research: From tank to bedside. Trends in neurosciences, 37(5), 264–278. 10.1016/j.tins.2014.02.011

Suriyampola, P. S., Shelton, D. S., Shukla, R., Roy, T., Bhat, A., & Martins, E. P. (2016). Zebrafish Social Behavior in the Wild. Zebrafish, 13(1), 1–8. 10.1089/zeb.2015.1159

Swaney, W. T., Ellwood, C., Davis, J. P., & Reddon, A. R. (2025). Familiarity preferences in zebrafish (Danio rerio) depend on shoal proximity. Journal of Fish Biology, 107(4), 1122–1128. 10.1111/jfb.15963

Tropepe, V., & Sive, H. L. (2003). Can zebrafish be used as a model to study the neurodevelopmental causes of autism? Genes, Brain, and Behavior, 2(5), 268–281. 10.1034/j.1601-183x.2003.00038.x

van Boxelaere, M., Clements, J., Callaerts, P., D’Hooge, R., & Callaerts-Vegh, Z. (2017). Unpredictable chronic mild stress differentially impairs social and contextual discrimination learning in two inbred mouse strains. PloS One, 12(11), e0188537. 10.1371/journal.pone.0188537

Varlinskaya, E. I., & Spear, L. P. (2012). Increases in anxiety-like behavior induced by acute stress are reversed by ethanol in adolescent but not adult rats. *Pharmacology*, Biochemistry, and Behavior, 100(3), 440–450. 10.1016/j.pbb.2011.10.010

Viechtbauer, W. (2010). Conducting Meta-Analyses in R with the metafor Package. Journal of Statistical Software, 36, 1–48. 10.18637/jss.v036.i03

Wei, D., Talwar, V., & Lin, D. (2021). Neural circuits of social behaviors: Innate yet flexible. Neuron, 109(10), 1600–1620. 10.1016/j.neuron.2021.02.012

Wright, D., Rimmer, L. B., Pritchard, V. L., Krause, J., & Butlin, R. K. (2003). Inter and intra-population variation in shoaling and boldness in the zebrafish (Danio rerio). Die Naturwissenschaften, 90(8), 374–377. 10.1007/s00114-003-0443-2

Wright, D., Ward, A. J. W., Croft, D. P., & Krause, J. (2006). Social organization, grouping, and domestication in fish. Zebrafish, 3(2), 141–155. 10.1089/zeb.2006.3.141

Zain, M. A., Pandy, V., Majeed, A. B. A., Wong, W. F., & Mohamed, Z. (2019). Chronic restraint stress impairs sociability butnot social recognition and spatial memoryin C57BL/6J mice. Experimental Animals, 68(1), 113–124. 10.1538/expanim.18-0078

