## Supplementary Figures for "Modulation of social behavior in adult zebrafish (*Danio rerio*): a systematic review and meta-analysis"

**Figure S1.** Funnel plots showing Hedges' g plotted against the standard error for social preference. Each point corresponds to one effect size.

**Figure S2.** Funnel plots showing Hedges' g plotted against the standard error for shoal cohesion. Each point corresponds to one effect size.

**Figure S3.** Funnel plots showing Hedges' g plotted against the standard error for anxiety-like behavior. Each point corresponds to one effect size.

### Figure S1

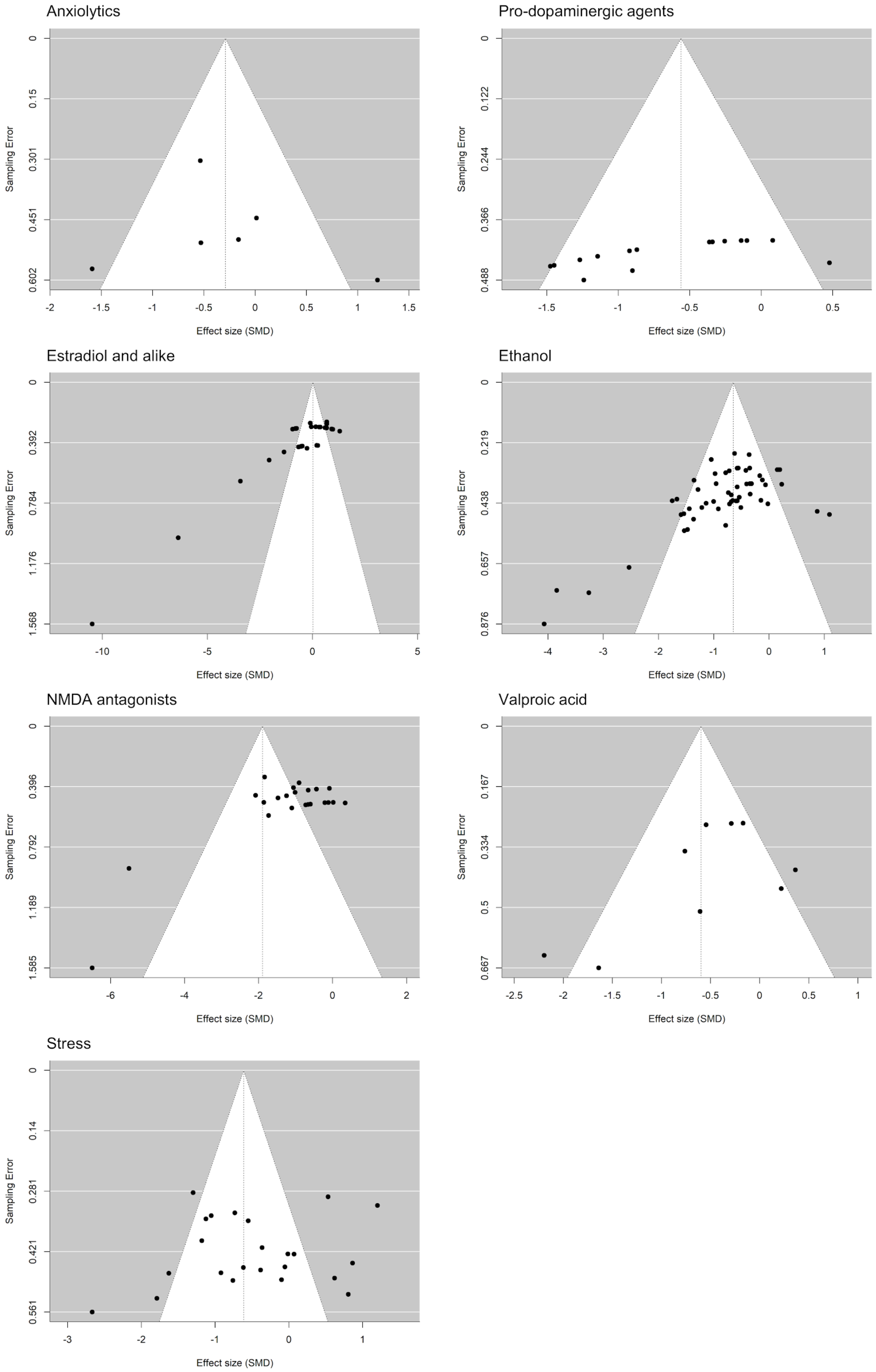
Anxiolytics
Sampling Error
0
0.15
0.301
0.451
0.602
-2
-1.5
-1
-0.5
0
0.5
1
1.5
Effect size (SMD)
Pro-dopaminergic agents
Sampling Error
0
0.122
0.244
0.366
0.488
-1.5
-1
-0.5
0
0.5
Effect size (SMD)
Estradiol and alike
Sampling Error
0
0.392
0.784
1.176
1.568
-10
-5
0
5
Effect size (SMD)
Ethanol
Sampling Error
0
0.219
0.438
0.657
0.876
-4
-3
-2
-1
0
1
Effect size (SMD)
NMDA antagonists
Sampling Error
0
0.396
0.792
1.189
1.585
-6
-4
-2
0
2
Effect size (SMD)
Valproic acid
Sampling Error
0
0.167
0.334
0.5
0.667
-2.5
-2
-1.5
-1
-0.5
0
0.5
1
Effect size (SMD)
Stress
Sampling Error
0
0.14
0.281
0.421
0.561
-3
-2
-1
0
1
Effect size (SMD)

### Figure S2

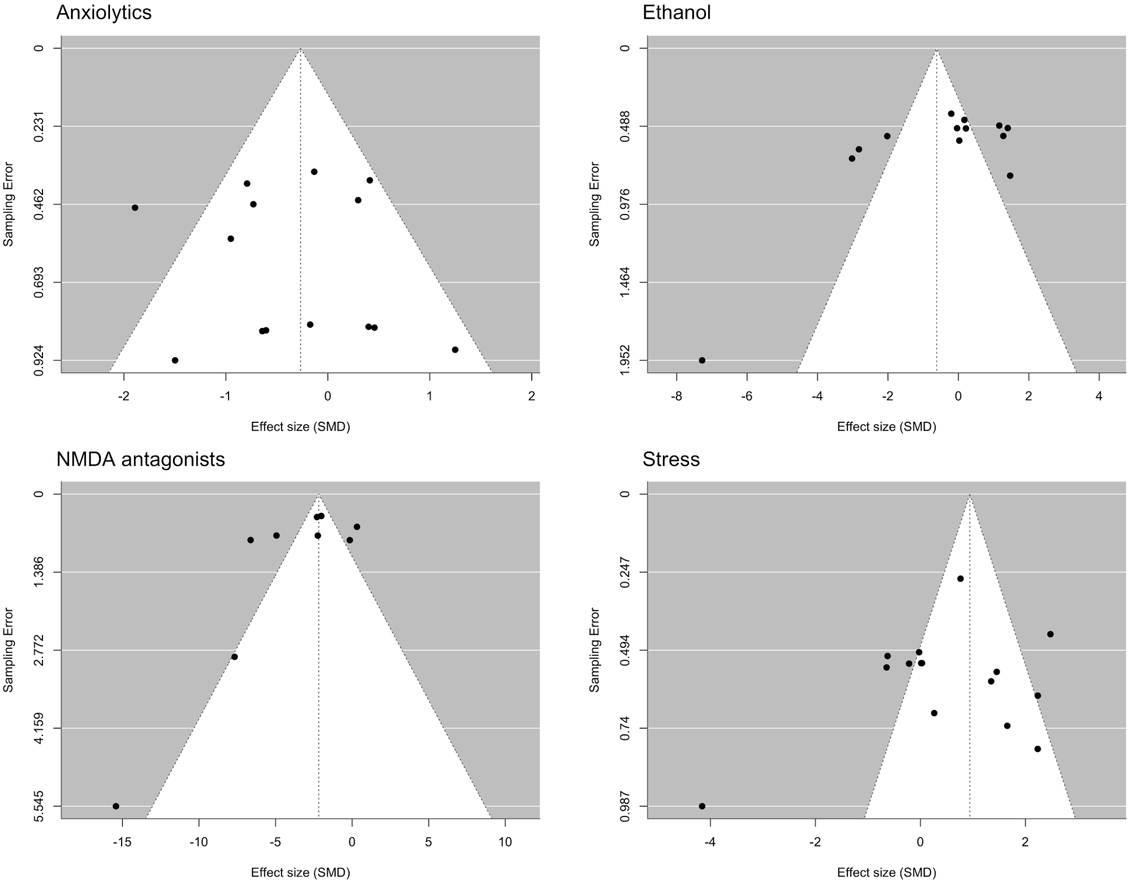
Anxiolytics
Ethanol
NMDA antagonists
Stress
Sampling Error
Effect size (SMD)

#### Figure S3

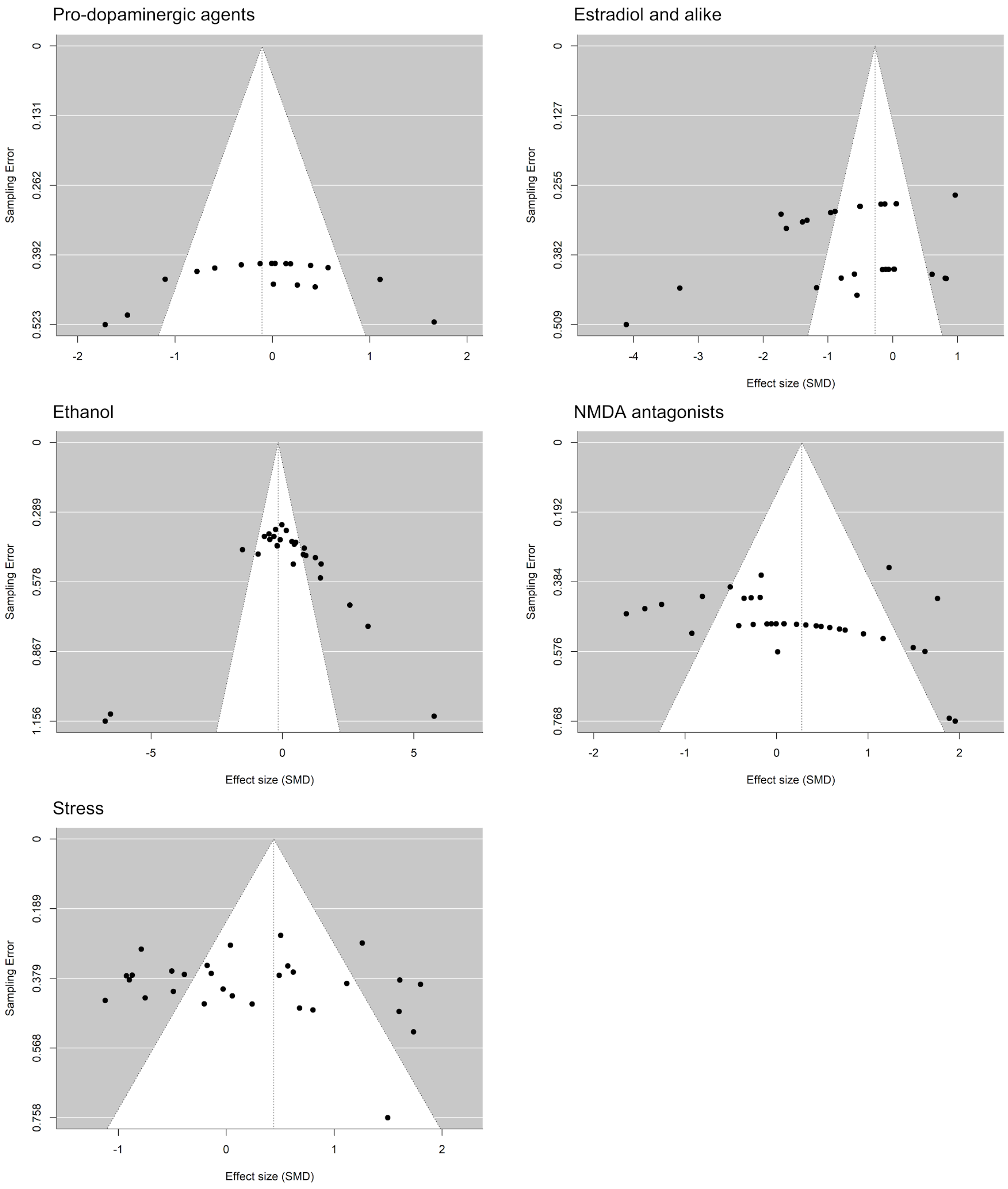
Pro-dopaminergic agents
Sampling Error
0
0.131
0.262
0.392
0.523
-2
-1
0
1
2
Estradiol and alike
Sampling Error
0
0.127
0.255
0.382
0.509
-4
-3
-2
-1
0
1
Effect size (SMD)
Ethanol
Sampling Error
0
0.289
0.578
0.867
1.156
-5
0
5
Effect size (SMD)
NMDA antagonists
Sampling Error
0
0.192
0.384
0.576
0.768
-2
-1
0
1
2
Effect size (SMD)
Stress
Sampling Error
0
0.189
0.379
0.568
0.758
-1
0
1
2
Effect size (SMD)
